# TMS-based neurofeedback training of mental finger individuation induces neuroplastic changes in the sensorimotor cortex

**DOI:** 10.1101/2024.05.16.594100

**Authors:** Ingrid Angela Odermatt, Manuel Schulthess-Lutz, Ernest Mihelj, Paige Howell, Caroline Heimhofer, Roisin McMackin, Kathy Ruddy, Patrick Freund, Sanne Kikkert, Nicole Wenderoth

## Abstract

Neurofeedback (NF) training based on motor imagery is increasingly used in neurorehabilitation with the aim to improve motor functions. However, the neuroplastic changes underpinning these improvements are poorly understood. Here, we used mental ‘finger individuation’, i.e., the selective facilitation of single finger representations without producing overt movements, as a model to study neuroplasticity induced by NF. To enhance mental finger individuation, we used transcranial magnetic stimulation (TMS)-based NF training. During motor imagery of individual finger movements, healthy participants were provided visual feedback on the size of motor evoked potentials, reflecting their finger-specific corticospinal excitability. We found that TMS-NF improved the top-down activation of finger-specific representations. First, intracortical inhibitory circuits in the primary motor cortex were tuned after training such that inhibition was selectively reduced for the finger that was mentally activated. Second, motor imagery finger representations in sensorimotor areas assessed with functional MRI became more distinct after training. Together, our results indicate that the neural underpinnings of finger individuation, a well-known model system for neuroplasticity, can be modified using TMS-NF guided motor imagery training. These findings demonstrate that TMS-NF induces neuroplasticity in the sensorimotor system, highlighting the promise of TMS-NF on the recovery of fine motor function.

## Introduction

Neural representations of individual body parts are activated when we execute movements and receive sensory inputs^1^. By now, it has been well established that these sensorimotor representations can also be activated without overt movement or sensory inputs, for example by attempted movements of completely paralysed^2,3^ or amputated body parts^4–6^, by motor planning that precedes motor execution^7^, or by motor imagery^8–10^, i.e., the pure mental simulation of movements^11^. Such activation of sensorimotor representations without motor execution can be used to control brain-computer interfaces (BCIs). BCIs detect and analyse brain signals and translate them into control commands to operate an external device (e.g., a prosthetic arm) or to neurofeedback (NF) that provides information about the current state of brain activity (referred to as BCI-NF). Repeatedly pairing the induced brain activity with NF allows users to gain volitional control of their brain activity and is thought to induce use-dependent neuroplasticity (for a review see^12,13^) which is the basis of restorative BCIs. Consequently, restorative BCIs are increasingly used in neurorehabilitation to aid motor recovery even in the absence of overt motor output, mostly following a stroke^14–16^, or spinal cord injury^17^. Such BCIs specifically aim to induce neuroplastic changes in sensorimotor pathways^12,18^. However, little is known about neuroplasticity induced by BCI-NF training beyond improvements in BCI-NF control itself^13,19^. Mixed results on the use of BCI-NF in stroke rehabilitation^20^ indicate that there is limited knowledge about the underlying neuroplastic changes of sensorimotor representations induced by a specific BCI-NF and how these neural changes might be reflected in improved motor performance following training.

Motor imagery of individual fingers targets sensorimotor finger representations that are well characterised. As such, mental ‘finger individuation’, i.e., the selective facilitation of single finger muscles without producing overt movements, can be used as a model to study neuroplasticity induced by BCI-NF. Importantly, the hallmarks of individuated finger movements can be assessed non-invasively using functional magnetic resonance imaging (fMRI) and transcranial magnetic stimulation (TMS). First, finger representations in the primary sensorimotor cortex (SM1) are somatotopically organised, providing a point-to-point correspondence of individual fingers to a specific area of the cortex^21,22^. Second, while these neural finger representations are largely overlapping, the individual activity patterns associated with individual fingers are separable in SM1^23,24^. Third, selectively moving individual fingers relies on a fine-tuned facilitation of the sensorimotor representations of a specific finger while inhibiting the others^25^. Specifically, intracortical circuits that regulate inhibition and facilitation of motoneurons within the primary motor cortex (M1) are involved in selective control of finger muscles during both motor execution^26,27^ and motor imagery^28–30^.

We previously developed a BCI-NF approach that enhances mental finger individuation through motor imagery^31^. In this BCI-NF training we use TMS to probe individual finger motor representations in M1 through motor imagery and provide visual feedback representing the TMS-induced motor evoked potentials (MEPs) of individual finger muscles as a read-out of corticospinal excitability. With this BCI-NF training, participants can learn to modulate their finger-specific corticospinal excitability^31^ (Fig. 1a).

**Figure 1.**
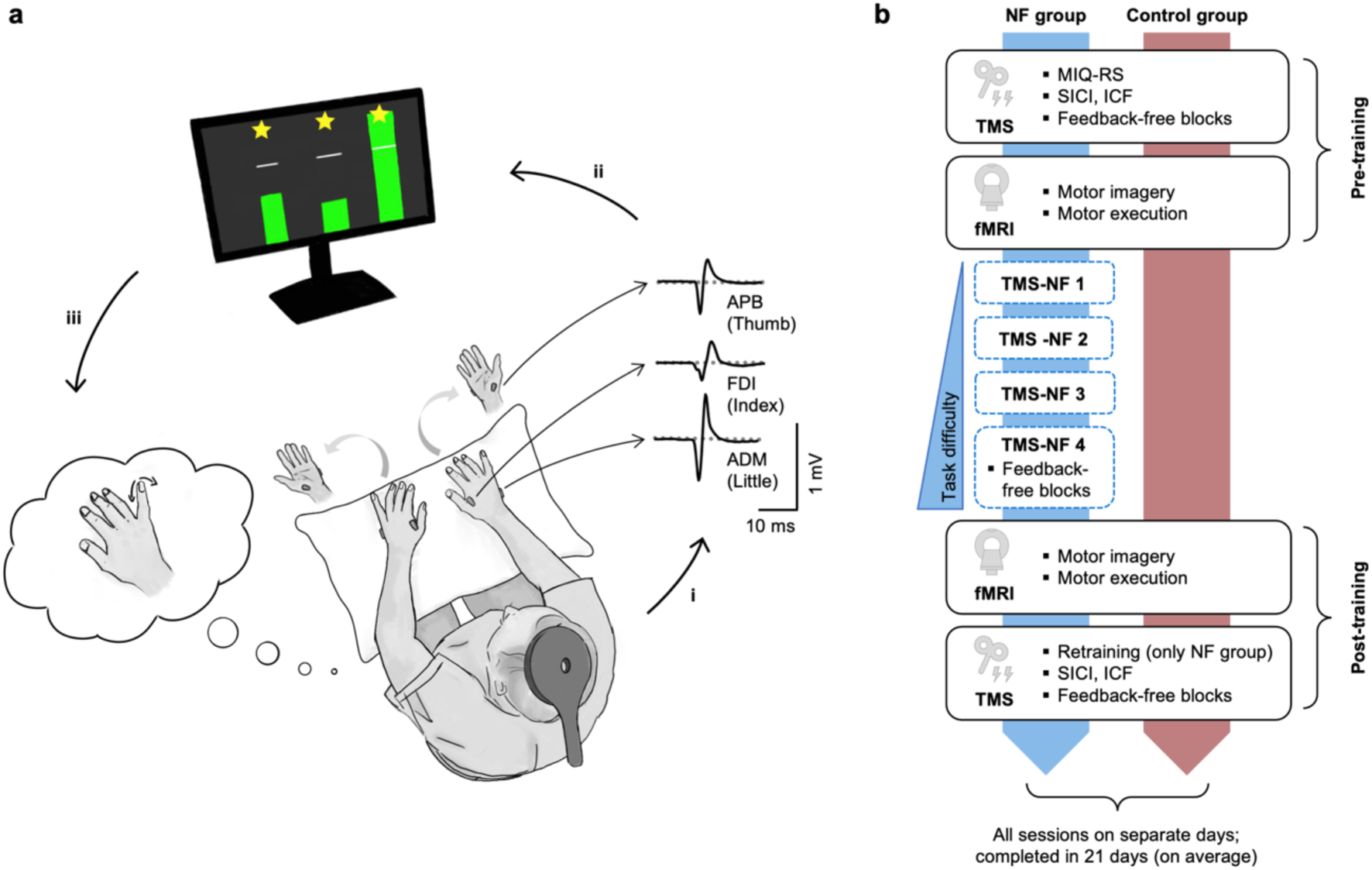
TMS-NF setup and study design**. a)** TMS-NF set-up. Participants sat in front of a computer screen and were instructed to imagine performing selective finger movements of the right hand (a little finger trial is visualised in the figure) while we recorded electromyography (EMG) of their finger muscles in both hands, i.e., left and right Abductor Pollicis Brevis (APB), First Dorsal Interosseus (FDI), and Abductor Digiti Minimi (ADM). i) During motor imagery, we applied a TMS pulse with a round coil to elicit motor evoked potentials (MEPs) simultaneously in the three right hand finger muscles. ii) We calculated the peak-to-peak amplitude of the MEPs, normalised them to the baseline (based on preceding rest trials), and displayed the normalised MEPs in the form of three bars (one for each finger muscle) as visual feedback on a screen. The white lines indicate no change from baseline, i.e., a normalised MEP of 1. If the bar of the instructed target finger was both above the white line and higher than the bars of the other two non-target fingers, the trial was deemed successful (green bars). Otherwise, it was deemed unsuccessful (red bars, not depicted here). In a successful trial, participants could earn up to three stars, one for each finger: The normalised MEP of the target finger had to be > 1.5; that of a non-target finger < 1. iii) Participants used the visual feedback to adapt their motor imagery strategies. **b)** Study design. The NF group (*n* = 16; blue) underwent four TMS-NF training sessions (TMS-NF 1-4) to train mental finger individuation. Task difficulty increased over sessions due to a transition from a blocked (i.e., one target finger per block) to an interleaved design (i.e., the target finger changed after each trial). The control group (*n* = 16; red) did not undergo any TMS-NF training. To measure the neural consequences of TMS-NF training, both groups underwent identical pre- and post-training TMS and fMRI sessions. During the first pre-training TMS session, we screened participants for their ability to perform kinaesthetic motor imagery using the Movement Imagery Questionnaire (MIQ-RS). In the pre- and post-training TMS sessions, we assessed short-interval intracortical inhibition (SICI) and intracortical facilitation (ICF) using paired-pulse TMS protocols. We further tested motor imagery performance in feedback-free blocks that were identical to the TMS-NF training blocks that had an interleaved trial order, but with occluded feedback. For the NF group, feedback-free blocks were also assessed at the end of the fourth TMS-NF training session. A short retraining period of TMS-NF was added to the start of the post-training TMS session for the NF group. In the pre- and post-training fMRI sessions, we measured brain activity during selective finger motor imagery and during the execution of a paced finger-tapping task.

Here, we used this TMS-NF approach to guide motor imagery to induce neuroplasticity. First, we aimed to understand the effects of TMS-NF training on neurophysiological mechanisms and whether intracortical circuits contribute to enhanced mental finger individuation in TMS-NF. We therefore used paired-pulse TMS protocols to probe short-interval intracortical inhibition (SICI) and intracortical facilitation (ICF) of M1 finger representations before and after TMS-NF training. We expected that a release of intracortical inhibition and an increase of facilitation for the target finger of motor imagery would be observed from pre- to post-training.

We then used fMRI and representational similarity analysis to examine whether improved finger individuation through TMS-NF training is related to more distinct, i.e., more separable, motor imagery finger representations after training. We further used a decoding analysis to investigate whether activity patterns elicited by imagined movements become more similar to those elicited by executed movements after TMS-NF training. Our main fMRI analysis focused on the SM1 hand cortex, as this brain region has been shown to exhibit high separability of finger representations^23,32^. We further explored changes in motor imagery finger representations in secondary motor areas, namely the ventral (PMv) and dorsal premotor cortex (PMd), and the supplementary motor area (SMA), as these areas have been implicated in motor imagery (for a review see^33,34^) and the encoding of imagined hand actions^9,10^.

## Results

We investigated the neural underpinnings of learning through motor imagery-based NF training. Specifically, we used TMS-NF training to enhance mental ‘finger individuation’, i.e., the selective facilitation of single finger muscles without producing overt movements (as in Mihelj et al.^31^): We instructed 16 participants to kinaesthetically imagine selective movements of the right thumb, index, or little finger. During motor imagery, we applied a TMS pulse over the contralateral M1 and computed the peak-to-peak amplitude of the TMS-evoked MEPs in the three right hand finger muscles (i.e., abductor pollicis brevis (APB), first dorsal interosseus (FDI), and abductor digiti minimi (ADM)). We then provided visual feedback representing MEP amplitudes normalised to rest (Fig. 1a). We trained participants in four TMS-NF sessions taking place on separate days. Over the training sessions, we gradually increased task difficulty by transitioning from a blocked to an interleaved trial order. All participants were able to successfully modulate corticospinal excitability for individual finger muscles in these training sessions (Supplementary Fig. 1a). We measured motor imagery performance pre and post TMS-NF training to quantify improvements in mental finger individuation. We further assessed plasticity of intracortical circuits in M1 induced by TMS-NF training using paired-pulse TMS protocols pre- and post-training. Finally, we assessed plasticity of neural finger representations in sensorimotor areas using fMRI pre- and post-training. A control group (*n* = 16) underwent identical pre and post measures as the NF group but did not undergo any TMS-NF training (Fig. 1b).

### TMS-NF training improves mental finger individuation

We first tested whether motor imagery performance changed after TMS-NF training. To do so, we assessed motor imagery performance pre- and post-training using a task identical to that used during TMS-NF training, but with occluded feedback. The trial order of these ‘feedback-free blocks’ was interleaved. We quantified motor imagery performance as the MEP target ratio, i.e., the ratio between the normalised MEP of the cued target finger muscle and the larger of the two non-target finger muscles normalised MEPs^31^. As such, an MEP target ratio greater than 1 indicates a finger-selective upregulation of corticospinal excitability.

The NF group improved motor imagery performance from pre- to post-training (*t*_(30.1)_ = -2.55, *p* = .02, Cohen’s *d* = 0.93, 95% CI for Cohen’s *d*: [ 0.17, 1.67]), whereas the control group did not (*t*_(29.6)_ = 0.57, *p* = .58, Cohen’s *d* = 0.20, 95% CI for Cohen’s *d*: [-0.52, 0.93]; significant Session (pre-training, post-training) by Group (NF, control) interaction: *F*_(1,30.89)_ = 4.69, *p* = .04, Cohen’s *d* = 0.78, 95% CI for Cohen’s *d*: [0.04, 1.50]; Fig. 2). In the pre-training session, there was no significant difference in motor imagery performance between the groups (*U* = 152, *p* = .38, *r*_b_ = .02, 95% CI for *r*_b_: [-0.21, 0.53]; BF_10_ = 0.42 indicating anecdotal evidence for the null hypothesis, i.e., no difference between the NF and the control group). During the TMS measurements, we strictly controlled for actual finger muscle activation (i.e., background EMG; bgEMG) by preventing a trial from proceeding if the bgEMG in any muscle exceeded 10 µV. Furthermore, we excluded all trials with bgEMG above 7 µV immediately before the TMS pulse in the offline analysis. Additionally, we controlled for potential effects of very subtle finger muscle activation by including the bgEMG target ratio as a covariate in the analysis reported above. Importantly, the bgEMG target ratio did not significantly contribute to the prediction of motor imagery performance (*F*_(1,48.31)_ = 0.51, *p* = .48, Cohen’s *d* = 0.21, 95% CI for Cohen’s *d*: [-0.36, 0.77]).

**Figure 2.**
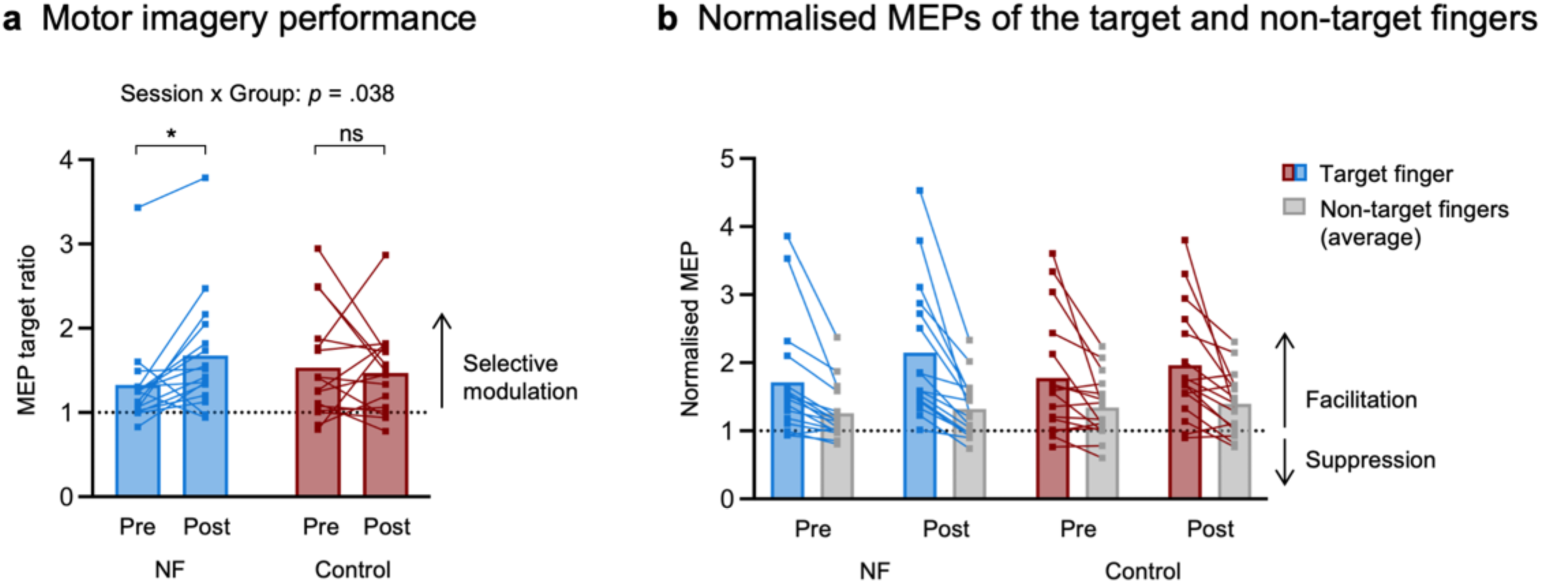
Motor imagery performance improves from pre to post TMS-NF training. **a)** MEP target ratio, i.e., the ratio between the normalised MEP (to the baseline at rest) of the target finger and the larger normalised MEP of the two non-target fingers. Values > 1 indicate a finger-selective modulation of corticospinal excitability. The data depicted corresponds to the feedback-free blocks acquired in the TMS pre- and post-training testing sessions for the NF and control groups. The MEP target ratio is a more conservative measure of finger-selective MEP modulation than comparing the MEPs of the target finger to the average of the non-target fingers as depicted in b). Therefore, statistical analysis was only performed on the MEP target ratio. **b**) Normalised MEPs of the target fingers (NF group = blue, control group = red) and the average normalised MEPs of the two non-target fingers (grey). This data is shown for visualisation merely. Squares depict data of individual participants. * *p* < .05; ns = non-significant.

These findings confirm that training with TMS-NF improved finger-selective modulation of corticospinal excitability. Importantly, they also demonstrate that these improvements in mental finger individuation translated to later sessions where participants did not receive any NF. This is a crucial precondition to interpret the neural changes that were assessed in the absence of NF.

### Intracortical inhibitory circuits are tuned following TMS-NF training

To investigate neural changes induced by TMS-NF training, we first tested for changes in neurophysiological circuits. As these intracortical circuits within M1 are highly relevant in shaping motor representations for skilled finger movements, we aimed to investigate the potential effects of TMS-NF training on two different circuits. Specifically, we used paired-pulse TMS protocols pre- and post-training to assess: (i) short-interval intracortical inhibition (SICI), which measures postsynaptic GABA_A_-ergic inhibition within M1^35,36^, and (ii) intracortical facilitation (ICF), which is thought to be dissociable from SICI circuits and to instead reflect glutamatergic facilitation^26,37^. We measured MEPs in the right index finger muscle (FDI) and assessed the two paired-pulse TMS protocols while participants imagined moving either the index finger or the thumb. This resulted in two motor imagery conditions where the index finger was either the target or a non-target finger. Here, we aimed to investigate if there was a release of SICI (and/or an increase of ICF) from pre- to post-training for a finger in the target condition relative to the non-target condition.

To test whether intracortical inhibition changed after training we calculated the pre- to post-training change in SICI. As such, positive scores indicate an increase in inhibition after TMS-NF training whereas negative scores indicate a decrease in inhibition after training. We then investigated whether these SICI change scores were different between the motor imagery conditions and between groups. In the NF group, we found that the change of SICI after training significantly differed for the target compared to the non-target condition. In other words, we observed a decrease in intracortical inhibition in the index finger if participants imagined moving the index finger compared to when they imagined to move the thumb (*t*_(30)_ = -2.39, *p* = 0.02, Cohen’s *d* = 0.85, 95% CI for Cohen’s *d*: [0.13, 1.56]), as opposed to the control group (no difference between conditions: *t*_(30)_ = 0.86, *p* = 0.39, Cohen’s *d* = 0.31, 95% CI for Cohen’s *d*: [-0.41, 1.02]; significant Motor imagery condition (target, non-target) by Group (NF, control) interaction (*F*_(1,30)_ = 5.29, *p* = 0.03, Cohen’s *d* = 0.84, 95% CI for Cohen’s *d*: [0.09, 1.58]; Fig. 3). This finding suggests that a release of intracortical inhibition for the mentally activated target finger representation may have enhanced the upregulation of the target finger’s MEP during motor imagery after TMS-NF training. Simultaneously, increased inhibition of non-target finger representations may have additionally contributed to the selectivity of the MEP modulation.

**Figure 3.**
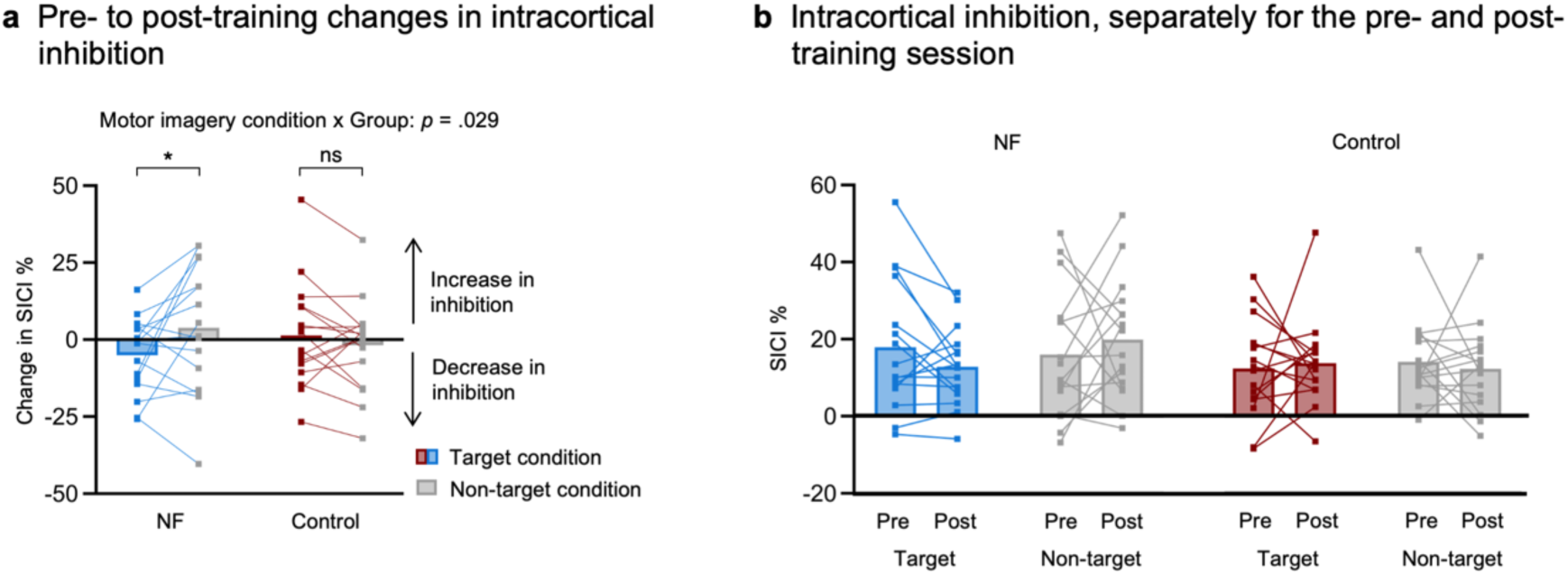
Intracortical inhibitory circuits are tuned after TMS-NF training. **a)** Pre- to post-training changes in short-interval intracortical inhibition (SICI). SICI was assessed with adaptive threshold hunting to determine the minimum testing stimulus intensity needed to elicit an MEP with an amplitude of at least 50% of the maximum MEP in 50% of trials. We measured SICI in the right index finger muscle during two motor imagery conditions: index as the target finger (motor imagery of index finger movements) vs index as an adjacent non-target finger (motor imagery of thumb movements). SICI is expressed as the % increase in the required testing stimulus intensity in the SICI protocol compared to a non-conditioned single pulse protocol during the same motor imagery condition. Positive scores indicate an increase in inhibition and negative scores indicate a decrease in inhibition after TMS-NF training. **b)** SICI for the pre- and post-training sessions separately. This data is shown for visualisation purposes only. Squares depict data of individual participants. * *p* < .05; ns = non-significant.

Analogous analyses were performed with ICF, but we did not find any significant effects of TMS-NF training (Supplementary Fig. 2).

### Single-finger motor imagery activates a fronto-parietal network

We first analysed univariate brain activity during motor imagery versus rest in the pre-training fMRI session. Our results confirmed that individual finger motor imagery (pre-training session, across all fingers and groups) activated a fronto-parietal network that is typically observed during motor imagery (for a review see ^33,34^; Fig. 4a). We observed activity in contralateral PMd and PMv with activity stretching into the M1 hand area, the inferior and superior parietal lobules, and bilateral SMA (see Supplementary Table 1a for a full list of activated clusters). We then computed univariate pre- to post-training changes in the activity levels during motor imagery. First, we tested whether these pre- to post-training changes in activity levels differed for the NF and the control groups. A whole-brain analysis did not reveal any significant group differences (see Supplementary Table 1b for pre- to post-training comparisons separately for both groups). Second, we investigated group differences in any activity level changes that predicted performance changes. We found that the largest significant cluster was located in M1 and overlapped with our main ROI encompassing the SM1 hand area (see Supplementary Table 1c for all significant clusters). For visualisation purposes and to ease interpretation, we then extracted the pre- to post-training change in activity levels under this M1 cluster per participant and correlated it with the corresponding MEP target ratio change (Fig. 4b). For the NF group, an increase in motor imagery performance was associated with a decrease in M1 activity (*r*_Pearson_ = -.75, *p* < .001, 95% CI: [- 0.91, -0.40]). For the control group, an increase in motor imagery performance was associated with an increase in M1 activity but this correlation did not reach significance (*r*_Spearman_ = .48, *p* = .06, 95% CI: [-0.02, 0.79]).

**Figure 4.**
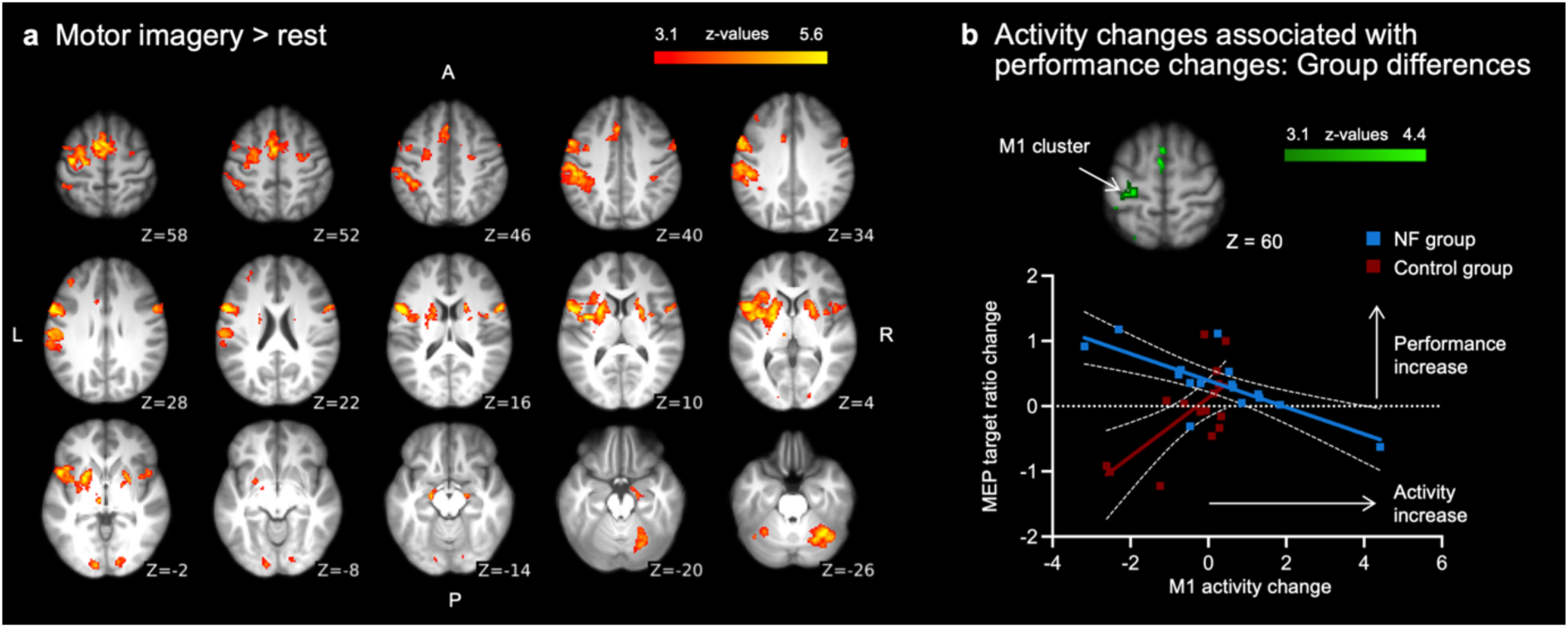
Motor imagery network and group differences of activity changes that predict performance changes. **a)** Whole-brain maps showing the overall activity during motor imagery (i.e., across all fingers and both groups) in the pre-training session. Single-finger motor imagery activated a fronto-parietal network and subcortical structures that are typical for motor imagery. **b)** Visualisation of the interaction effect in the M1 cluster demonstrating that the relationship between pre- to post-training changes in M1 activity and motor imagery performance changes differs between the NF and control groups. Changes in activity level (z-values) are depicted on the x-axis, with positive values showing an increase in activity from pre- to post-training. Changes in motor imagery performance are depicted on the y-axis, with positive values indicating an improvement from pre- to post-training. Squares depict data of individual participants, coloured lines show the best fit, and white dotted lines show the 95% confidence bands.

### Neural finger representations activated by motor imagery become more distinct following TMS-NF training

Next, we performed an in-depth investigation of plasticity of finger representations in SM1 and an exploratory analysis of plasticity of finger representations in secondary motor areas (Fig. 5a). We expected a co-involvement of M1 and the primary somatosensory cortex (S1) during motor imagery, with M1 being implicated in MEP modulation^38,39^ and S1 containing the imagined sensory consequences of imagined movements^40,41^. Specifically, we used multivariate pattern analysis (MVPA) to study changes in fine-grained finger representations induced by TMS-NF training. MVPA allows to investigate the intricate relationship between experimental conditions and activity patterns across voxels, which is particularly advantageous in the case of overlapping (finger) representations as in SM1^23,24,32^. With representational similarity analysis (RSA) we examined the relationship between activity patterns elicited by imagined finger movements in an anatomically defined ROI, and then averaged the resulting inter-finger distances across finger pairs within each participant to estimate the average inter-finger separability (or finger representation strength). We expected that after TMS-NF training individuated finger motor imagery would elicit activity patterns in SM1 that contain increased information content to distinguish between fingers. If motor imagery finger representations would become more distinct across fingers, then the separability would increase.

**Figure 5.**
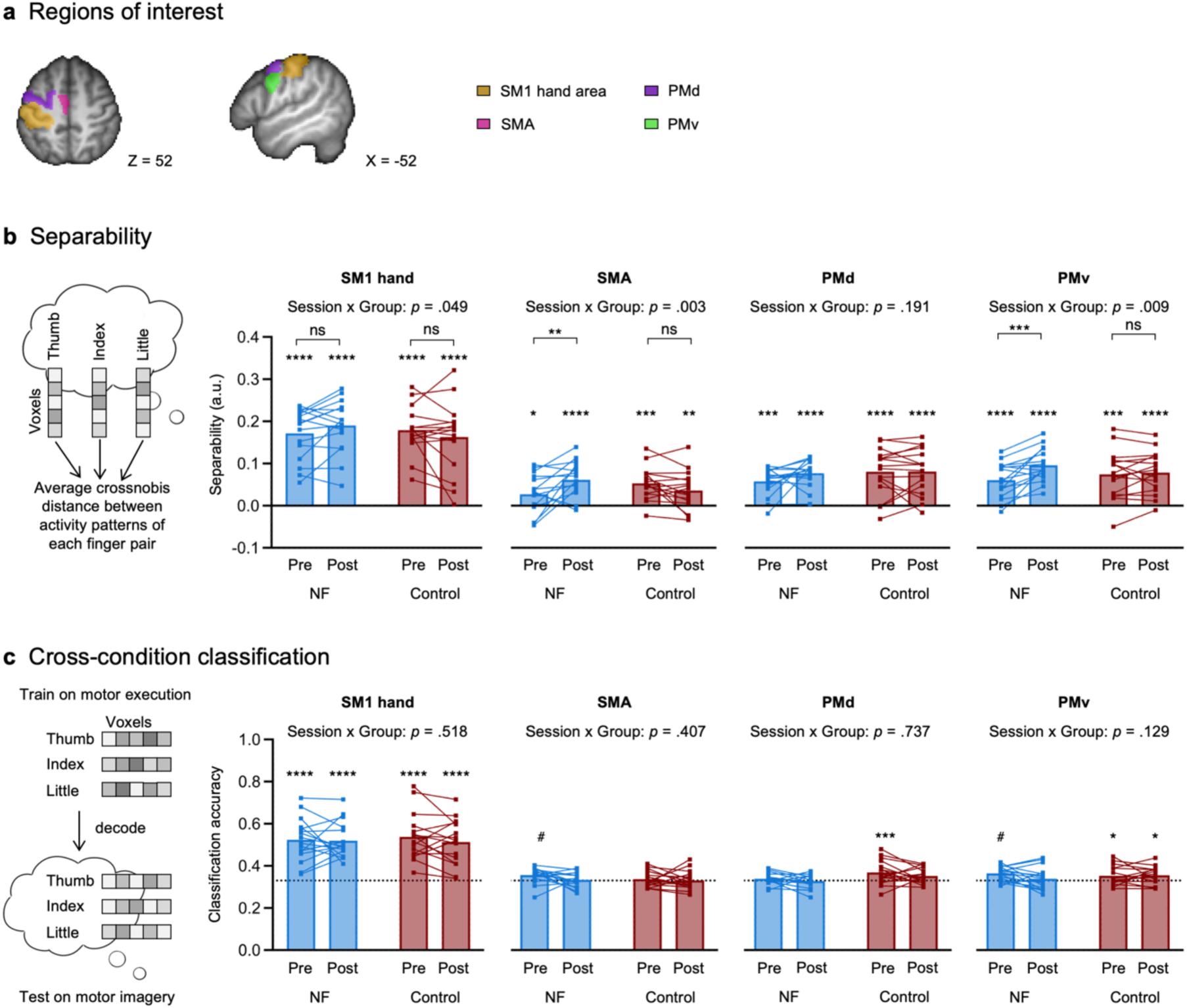
Finger representations activated by single-finger motor imagery become more separable following TMS-NF training, but do not become more similar to motor execution. **a)** Anatomically defined regions of interest (ROIs) used for multivariate pattern analysis. **b)** Finger separability, measured as the average inter-finger distance, of the representational structure of imagined finger movements in the SM1 hand, SMA, PMd and PMv ROIs for the NF and control groups. The distance is computed as the average cross-validated Mahalanobis (crossnobis) distance between activity patterns elicited by single-finger motor imagery of each finger pair. Asterisks on top of the bars indicate significant differences from 0 (FDR-corrected within each ROI and group). **c)** Cross-condition classification accuracy. A linear support vector machine was trained separately for each participant on all motor execution trials across both the pre- and post-training sessions to predict the motor imagery trials in the pre- and post-training sessions. The dotted line represents the empirical chance level (33.33%). Asterisks refer to the statistical difference of classification accuracy from the empirical chance level (FDR-corrected within each ROI and group). Squares depict data of individual participants. **** p < .0001; *** p < .001; ** p < .01; * p < .05; # p < .10; ns = non-significant.

Inter-finger separability was greater than 0 in all ROIs for all measured time points and groups (all *p*_(FDR)_ < .033), indicating that the activity patterns in SM1 and all tested secondary motor areas contained finger-specific information. We found that finger representations activated by motor imagery became more separable in SM1 following TMS-NF training for the NF group compared to the control group (significant Session by Group interaction; *F*_(1,30)_ = 4.22, *p* = .049, Cohen’s *d* = 0.75, 95% CI for Cohen’s *d*: [0.00, 1.48]; Fig. 5b). However, post-hoc contrasts comparing the pre- to post-training sessions separately for the groups, did not yield significant differences (NF group: *t*_(30)_ = -1.56, *p* = .13, Cohen’s *d* = 0.55, 95% CI for Cohen’s *d*: [-0.17, 1.28]; control group: *t*_(30)_ = 1.34, *p* = .19, Cohen’s *d* = 0.47, 95% CI for Cohen’s *d*: [-0.25, 1.20]). In secondary motor areas, we found significant Session by Group interactions for SMA (*F*_(1,30)_ = 10.56, *p* = .003, Cohen’s *d* = 1.19, 95% CI for Cohen’s *d*: [0.40, 1.95]), and PMv (*F*_(1,30)_ = 7.74, *p* = .009, Cohen’s *d* = 1.02, 95% CI for Cohen’s *d*: [0.25, 1.77]), but not for PMd (*F*_(1,30)_ = 1.79, *p* = .19, Cohen’s *d* = 0.49, 95% CI for Cohen’s *d*: [-0.24, 1.21]). Separability in SMA (*t*_(30)_ = -3.07, *p* = .005, Cohen’s *d* = 1.09, 95% CI for Cohen’s *d*: [0.35, 1.82]) and PMv (*t*_(30)_ = - 4.48, *p* = .0001, Cohen’s *d* = 1.58, 95% CI for Cohen’s *d*: [0.81, 2.36]) increased significantly from pre- to post-training for the NF group but not for the control group (SMA: *t*_(30)_ = 1.53, *p* = .14, Cohen’s *d* = 0.54, 95% CI for Cohen’s *d*: [-0.18, 1.26]; PMv: *t*_(30)_ = -0.54, *p* = .59, Cohen’s *d* = 0.19, 95% CI for Cohen’s *d*: [-0.52, 0.91]).

### Activity patterns elicited during individual finger motor imagery do not become more similar to those observed during motor execution after TMS -NF training

To investigate whether neural activity patterns elicited by individual finger motor imagery became more similar to those observed during motor execution following TMS-NF training, we performed a cross-condition decoding analysis. Specifically, we trained a linear support vector machine to decode fingers during the motor execution task (i.e. paced individual finger tapping; Supplementary Fig. 3) and tested whether this decoder could be generalised to the motor imagery task, i.e., across another condition. If there is shared information in the activity patterns elicited by imagined and executed finger movements in a given ROI, then this would be reflected in a cross-condition classification accuracy above chance level. If the activity patterns elicited by motor imagery would become more similar to motor execution after TMS-NF training, then the cross-condition classification would increase from pre- to post-training.

We found consistent classification accuracies greater than chance for all sessions and groups for the SM1 hand area but not for secondary motor areas (Fig. 5c). However, the cross-condition classification accuracy in the SM1 hand area did not significantly differ across sessions or groups (no significant main effects and no Group by Session interaction: *F*_(1,30)_ = 0.43, *p* = .52, Cohen’s *d* = 0.24, 95% CI for Cohen’s *d*: [-0.48, 0.96]). Bayesian tests provided moderate evidence for the null hypothesis, i.e., no change from pre- to post-training sessions for the NF group (BF_10_ = 0.26) and anecdotal evidence for the null hypothesis for the control group (BF_10_ = 0.50).

### 2.5 Neural changes do not directly predict changes in motor imagery performance

Finally, we explored whether the improved motor imagery performance in the NF group (i.e., pre- to post-training change in MEP target ratio) related to our main neural outcome measures (i.e., SICI % changes in the target condition, separability changes in the SM1 hand area, and cross-condition classification accuracy changes in the SM1 hand area). A multiple linear regression revealed that the changes in the measured neural mechanisms paralleled an improvement in motor imagery performance in the NF group but did not directly predict the observed changes (multiple *r*^2^ = .12; separability: *t* = 1.13, *p* = .28; cross-condition classification: *t* = -0.90, *p* = .39; SICI %: *t* = 0.18, *p* = .86). Similar results were found when including both groups in the analysis (i.e., NF and control groups; multiple *r*^2^ = .05; separability: *t* = 0.43, *p* = .67; cross-condition classification: *t* = 0.52, *p* = .61; SICI %: *t* = 0.71, *p* = 49).

## Discussion

In this study, we investigated neuroplastic changes induced by mental finger individuation training that was guided by TMS-NF. Replicating our previous work^31^, we found that TMS-NF training enabled participants to selectively upregulate corticospinal excitability of a target finger while simultaneously downregulating corticospinal excitability of other finger representations. Our new findings demonstrate that this finger-specific training effect is mediated by tuning inhibitory circuits in M1: After TMS-NF training, GABA_A_-ergic inhibition was released if a finger was targeted, while inhibition was increased when the finger was not targeted. We further found that through TMS-NF training, activity patterns underpinning single-finger motor imagery became more distinct in SM1, SMA, and PMv. Together, our findings demonstrate that TMS-NF to guide motor imagery training induces neuroplastic changes that go beyond test-retest effects.

Using neurophysiological assessments, we demonstrated that following TMS-NF training, the selective activation of an M1 finger representation through motor imagery was associated with a release of intracortical inhibition measured in this target finger muscle. Relative to that, when measuring in that same finger muscle during motor imagery of another finger, there was an increase in intracortical inhibition. Intracortical inhibition, assessed with SICI, is thought to be driven by inhibitory interneurons in M1^42^ that are crucial for the fine-tuned activation and suppression of motor representations^26,27^. Our results align with a previous BCI-NF study that showed a decrease in intracortical inhibition for the agonist (or target) muscle compared to rest while it remained unaffected for an antagonist (or non-target) muscle during motor imagery of wrist movements^43^. In line with this study^43^, we did not find changes in glutamatergic facilitatory circuits after TMS-training. Even after physical training, no clear training effects on ICF have been reported^26^. These studies and our results suggest that disinhibition (e.g., by a release of SICI) rather than facilitation might be essential for intracortical plastic changes^26^. During BCI-NF training, a release of SICI might not induce a general increase of corticospinal excitability but rather modulate the specific activation of adjacent sensorimotor representations^43^. This modulation is thought to be driven by tuning horizontal connections, similar to the corticospinal excitability changes that are observed during executed movements^43^. Our results corroborate this finding by showing similar modulatory effects of SICI during mental activation of neighbouring finger muscle representations in a feedback-free scenario after TMS-NF training. These findings mirror changes in SICI during execution of individual finger movements and demonstrate that modulation of SICI might enhance the selectivity of finger muscle activation in M1^27^. In our study, the modulatory effects on SICI during mental finger individuation suggest that tuning of intracortical inhibitory mechanisms may have ‘shaped’ motor imagery finger representations during TMS-NF training.

The finding that the TMS-NF group learned to selectively activate single-finger representations is further supported by our fMRI results. Previous fMRI studies have shown that individual SM1 finger representations can be activated through top-down processes, i.e., without overt movements or sensory stimulation, such as through attempted^3–5^ and planned^7^ movements or observed touch^44^. Here, we add to that growing body of literature by demonstrating that motor imagery of individual fingers evoked separable activity patterns in SM1 across all participants in both fMRI sessions. Importantly, following TMS-NF training, these SM1 representations became more finger-specific (i.e., inter-finger distances increased from pre- to post-training) compared to the control group that did not undergo any TMS-NF training. To our knowledge, our study is the first to show fMRI changes in pure top-down activated SM1 finger representations following motor imagery based BCI-NF training. This could reflect a more selective activation of single finger representations following training due to less enslavement, i.e., less activation of non-target fingers. This may parallel effects found in motor execution, where more enslavement has been associated with more overlapping finger representations^23,24^.

Most studies that investigated plasticity of SM1 finger representations used motor execution paradigms to elicit finger-specific activity patterns. These SM1 finger representations activated by executed movements have been shown to be relatively stable over time^45^ and training interventions^46^. Specifically, five weeks of training to perform specific finger movement sequences, finger movement representations did not change in S1 or M1^46^. Even after life-long expert-learning^32,47,48^ or drastic changes in sensorimotor experiences^3–5^ finger movement representations generally remained stable or only underwent subtle changes. For example, S1 finger representations activated by phantom finger movements of amputees’ missing hand or tetraplegic patients’ paralysed hand showed similar finger somatotopy as healthy controls^3–5,49^. However, few studies have demonstrated changes in finger representations after long-term learning or certain interventions. It was shown that finger movement representations had increased overlap in professional compared to amateur musicians in M1 (but not S1)^32^. Additionally, gluing fingers together for 24h^50^ or blocking nerves in specific fingers for approximately 5h altered S1 finger movement representations^51^. The majority of studies however reported stable representations. In contrast, our findings show changes in motor imagery finger representations in SM1 following TMS-NF training. It is possible that top-down activated finger representations are more malleable compared to finger movement representations. In line with that, a study combined motor execution with mental strategies and showed that manipulated online fMRI-NF of finger representations can teach individuals to volitionally shift SM1 representations of some fingers during individuated finger movements^52^.

We found that without any training intervention (i.e., in the control group; test-retest effects) separability of finger representations was slightly decreased in the second testing sessions, although this difference did not reach significance. However, relative to this slight decrease, we observed an increase in separability for the NF group, resulting in a significant Session by Group interaction. The observed trend towards a slight decrease in separability without TMS-NF is in line with our previous study that combined TMS-NF during mental finger individuation with electroencephalography (EEG)^31^: Mihelj et al. included a control group which performed motor imagery but did not receive veridical feedback. Separability scores were calculated from EEG and revealed a slight decrease for the control group over training sessions, while there was an increase for the TMS-NF group^31^.

We found that a decrease of M1 activity during motor imagery was associated with better motor imagery performance in the NF group, while increased separability of SM1 finger representations did not directly correlate with the performance changes. These results suggest that with better performance following TMS-NF training the activity level in M1 decreased while the information content to distinguish between fingers remained stable. This might reflect a more efficient, i.e., more targeted activation of finger representations. Previous research has shown that more accentuated sensory representations were accompanied with lower activity levels^53^. However, it is important to note that the lack of a significant correlation between SM1 finger representation separability and performance changes should be interpreted with caution. The lack of a significant correlation does not necessarily indicate that there is no relationship, it may instead be explained by a lack of statistical power^54^.

At the whole brain level, executed and imagined movements have been shown to predominately activate the same network of areas^33,34,55^. Their neural representations are thought to share a low-to-moderate degree of similarity^10^. In line with this, we demonstrated that a decoder trained on SM1 activity patterns elicited by executed finger movements successfully generalised to imagined finger movements. However, we did not find that the resemblance of activity patterns elicited by imagined and executed finger movements differed between groups or changed due to training. This suggests that motor imagery finger representations did not become more similar to motor execution finger representations through training. Although the shared neural code of finger representations elicited by motor imagery and motor execution could still be detected in SM1, it is possible that task differences might have masked an increased resemblance of motor imagery and motor execution representations induced by TMS-NF training. We did not restrict participants to perform specific imagined movements but instead allowed them to find and develop their own motor imagery strategies during TMS-NF training. As a result, strategies used during motor imagery varied from, for example, button pressing, making circles with the cued finger, touching a surface, to finger abduction (see Supplementary Table 2c for self-reported motor imagery strategies during the fMRI sessions). Movement execution by contrast consisted of a paced button press task. However, it is also possible that motor imagery and execution rely on different neural substrates within M1, with motor imagery being represented in superficial rather than the deep layers, while motor execution was represented in both superficial and deep layers^56^. Our fMRI approach did not allow us to investigate such potential layer-specific effects.

What processes may have driven the neuroplastic changes in SM1 induced by motor imagery combined with TMS-NF? We suggest that the observed effects on intracortical inhibition and separability of top-down finger representations may have been caused by an interplay of multiple processes^19^. First, use-dependent plasticity in SM1 has been frequently demonstrated for motor execution^57,58^ and motor imagery tasks^59–61^ and it is likely that this mechanism, possibly driven by long-term potentiation (LTP)-like plasticity, has been triggered by repeated practice with TMS-NF^62^. Second, gaining control of BCI-NF via motor imagery may additionally reflect skill learning that involves a network beyond SM1. It is therefore possible that any changes in SM1 representations may emerge due to interconnections with various other, higher-order, brain areas, such as premotor and parietal association areas. Indeed, studies investigating effective connectivity during motor imagery suggest that SMA, PMv, and PMd are bidirectionally connected to each other and to SM1^63–65^. In line with this, we observed higher separability of motor imagery finger representations in SMA and PMv following TMS-NF training. Previous work indicates that controlling BCI-NF via motor imagery is a skill that, once acquired, can be maintained over long periods without training^66,67^, further supporting that skill learning may be involved in BCI-NF training. Likely, an interplay of inter- and intrahemispherically^68^ connected areas in the sensorimotor network has contributed to the effects we found in SM1. Finally, studies have shown that it is possible to activate somatotopic S1 hand representations by merely directing attention to individual fingers^69,70^. It is therefore possible that through improving attentional processes, participants might have targeted motor imagery representations more selectively. Importantly, these possible mechanisms are not mutually exclusive, and it is likely that neuroplastic, skill learning dependent, and attentional processes contributed to the observed changes in SM1 finger representations following TMS-NF training.

The neural changes induced by TMS-NF training demonstrate the promise of TMS-NF for use in a clinical setting as a BCI-NF training to restore fine motor control. This is further supported by the high aptitude rate and the rapid learning reported in TMS-NF studies if participants receive informative feedback^31,66,71–73^. Additionally, we observed a translation of improved performance during the training to a feedback-free scenario after training. Once the motor imagery strategies were acquired, 14 out of 16 participants were able to apply their strategies to reach an improved motor imagery performance without receiving NF. This finding is in line with our previous work using a simplified TMS-NF set-up in which participants were able to maintain performance in a feedback-free scenario even six months after training ^48^. Regaining hand functions has been reported as one of the most important therapy goals by tetraplegic and stroke patients^74^. TMS-NF might offer a rehabilitation strategy that can be employed already in the early stages after for example a stroke or a spinal cord injury when patients are not yet able to engage in physical training. The simplified TMS-NF setup has previously been tested in a clinical setting. In a feasibility study, subacute stroke patients (*n* = 7) who received TMS-NF learned over four training sessions to increase corticospinal excitability in paretic muscles^75^. Larger trials with more participants and longer training periods to test for the effects of TMS-NF on functional motor recovery will give further insight into its clinical relevance. Importantly, our findings also open new avenues to investigate the extension of TMS-NF as a tool to shape top-down sensorimotor representations. Such training could improve control in other BCIs that rely on clearly separable neural activity patterns or be beneficial in neurological disorders associated with aberrant or disorganised sensorimotor representations.

In summary, our results show that TMS-NF improved the top-down activation of finger-specific motor representations by tuning intracortical inhibitory networks in M1 such that inhibition was selectively reduced for a finger that was mentally activated while it was increased for another finger. These neurophysiological findings were further corroborated by fMRI revealing that finger representation became more distinct after training consistent with a sharper, less overlapping recruitment of the neural populations representing a specific finger. Together, our results indicate that the neural underpinnings of finger individuation, a well-known model system for neuroplasticity, can be modified using motor imagery training that is guided by TMS-NF. With this proof-of-principle study we demonstrate that BCI-NF training can indeed promote neuroplasticity that may be relevant for motor recovery.

## Material and Methods

### Participants

For this study, we recruited 46 participants. Inclusion criteria were: No use of medication acting on the central nervous system, no neurological and psychiatric disorders, right-handed according to the Edinburgh Handedness Inventory^76^, normal or corrected-to-normal vision, and no TMS^77,78^ and MRI contraindications. At the start of the study onset (i.e., at the beginning of the pre-training TMS session), we screened participants for their ability to perform kinaesthetic motor imagery using the kinaesthetic subscale of the Movement Imagery Questionnaire – Revised second version (MIQ-RS^79,80^). In this questionnaire, participants are instructed to perform and then kinaesthetically imagine movements and rate this mental task from 1 (very hard to feel) to 7 (very easy to feel). We asked participants with low scores, i.e., more than 1 SD below the mean score reported in Gregg et al.^79^, whether they were able to mentally simulate the kinaesthetic experience of movements. If participants negated, we excluded them from the study.

We excluded a total of 14 participants after study enrolment due to: (i) reported difficulty to perform kinaesthetic motor imagery (2 participants), (ii) a high resting motor threshold (RMT) that was above 80% of the maximum stimulator output (MSO) and resulted in difficulties to find a suitable testing intensity (6 participants), (iii) reported discomfort during TMS or fMRI (3 participants), persistent background electromyography amplitude (bgEMG) that exceeded the online bgEMG control ( >10 μV) during the first TMS session (1 participant), (iv) excessive head motion in the first fMRI session, i.e., a mean displacement >1.1mm (corresponding to half a voxel size) in the majority of runs (1 participant), or (v) being unsure about MRI contraindications (1 participant). Testing was completed by 16 participants in the neurofeedback group (NF; age (mean ± SD): 25.1 ± 2.8 years; 8 females) and 16 participants in the control group (age: 26.4 ± 2.7 years; 8 females), adhering to the sample size calculation that was made prior to study onset (using G*Power v3.1, based on the effect size reported in Mihelj et al.^31^). The participants who completed testing did not report any major side effects after the TMS sessions. All research procedures were approved by the Cantonal Ethics Committee Zurich (BASEC Nr. 2018-01078) and were conducted in accordance with the declaration of Helsinki. All participants provided written informed consent prior to study onset.

### Experimental procedure

The NF group underwent four sessions of TMS-NF to train individuation of imagined finger movements. Additionally, we conducted pre- and post-training TMS and fMRI testing sessions to measure the neural consequences of TMS-NF (Fig. 1b). In the pre- and post-training TMS sessions we used paired-pulse TMS protocols to quantify effects of TMS-NF on inhibition and facilitation in the primary motor cortex (M1) during motor imagery. In the pre- and post-training fMRI sessions, we acquired brain activity during imagined and executed selective finger movements to investigate neural finger representations. During the pre- and post-training TMS sessions we additionally assessed motor imagery performance in feedback-free blocks, i.e., identical to TMS-NF, but with occluded feedback. We also assessed such feedback-free blocks at the end of the fourth (and last) TMS-NF session for the NF group. This allowed us to investigate the stability of motor imagery performance by comparing the measurement directly after TMS-NF training to the measurement in the post-training TMS session. Note that for the first three participants we assessed the feedback-free blocks at the start of the and the end of the fourth TMS-NF session rather than in the pre- and post-training TMS sessions. The control group did not receive any TMS-NF training but underwent identical pre- and post-training sessions as the NF group to control for test-retest effects. Importantly, we have already shown that a control group that received uninformative NF did not improve their ability to up-vs downregulate (finger-selective) modulation of MEPs^31,66^. For one participant of the NF group, we repeated the post-training TMS session due to technical issues.

In the pre-training sessions, the NF and the control group received identical, standardized instructions to imagine selective movements with the cued finger and were provided example strategies based on Mihelj et al.^31^ and Ruddy et al.^66^ (see Supplementary Table 2a for verbatim instructions, and Supplementary Table 2b and 2c for self-reported strategies). For the post-training sessions, we instructed the NF group to apply the motor imagery strategies that they had acquired during the TMS-NF training.

We kept the experimenter and time of the day for the testing and training sessions consistent within each participant. All sessions took place on separate days and the whole study was completed in an average of 21 days (NF group (mean ± SD): 19.5 ± 5.5; control group (mean ± SD): 21.7 ± 13.9).

### TMS and EMG setup

During the TMS sessions participants sat in a comfortable chair with a headrest and placed their arms on a pillow on their lap. Surface EMG (Trigno Wireless, Delsys) was recorded from the left and right thumb (Abductor Pollicis Brevis; APB), index finger (First Dorsal Interosseus; FDI), and little finger (Abductor Digiti Minimi; ADM). EMG data were sampled at 1926 Hz (National Instruments, Austin, Texas), amplified, and stored on a PC for offline analysis. For TMS-NF, a round coil with a 90 mm loop diameter was connected to a Magstim 200 stimulator (Magstim, Whitland, UK) to deliver single-pulse monophasic TMS. We used a round coil for TMS-NF to achieve a less focal stimulation. As such, we were able to elicit motor evoked potentials (MEPs) in all three measured finger muscles of the right hand in the same coil position as in the setup of Mihelj et al.^31^. For paired-pulse TMS protocols, a 70 mm figure-of-eight coil was connected to two coupled Magstim stimulators. Here, we used a coil to allow for a more focal stimulation and optimally target the M1 representation of the right FDI. All stimuli were provided using custom MATLAB scripts (MATLAB 2020b, MathWorks) and Psychophysics Toolbox-3^81,82^.

### TMS-based neurofeedback task

We used similar procedures as in Mihelj et al.^31^ to train participants to selectively modulate their corticospinal excitability through motor imagery using TMS-NF. A TMS-NF trial started with a preparatory rest period of 1-2 s. During this time, the bgEMG of all measured finger muscles on the left and right hand was computed as the root mean square (rms) of the EMG signal within a sliding window of 100 ms. Participants saw six dots on the screen, representing the bgEMG of the individual muscles. The dots were green when the bgEMG was < 10 μV and turned red otherwise. Only when the bgEMG in all muscles was < 10 μV for a minimum of 1 s did the trial proceed to the motor imagery (or rest) period. During this period a visual cue appeared on the screen that instructed the participant to perform finger-selective motor imagery of the right hand (‘thumb’, ‘index’, or ‘little’) or to rest (‘rest’). The first ten trials in each block were rest trials, which we collected to determine a baseline for each finger muscle. The motor imagery (or rest) period of a trial lasted for a jittered period of 4-6 s to avoid anticipation effects for the TMS pulse^83^. If the bgEMG rms exceeded 10 μV in any muscle during this period, the TMS pulse was only sent once the bgEMG was below the threshold for the predefined motor imagery duration. The aim of the bgEMG control was to prevent participants from making subtle movements or muscle contractions as to ensure that any MEP modulation was caused solely by motor imagery. The bgEMG control only stopped in the last 0.5 s before the TMS pulse was applied. The dots remained green during this period, regardless of the bgEMG values. After each TMS pulse, we computed the MEP peak-to-peak amplitudes of the three right-hand finger muscles. The feedback (or fixation cross for rest trials) was displayed 1 s after the TMS pulse and lasted 3 s. The normalised MEP amplitudes were computed by dividing the MEP amplitude of a finger muscle by the rest MEP amplitude of the same finger muscle. This rest MEP amplitude was based on nine rest trials of the corresponding block, disregarding the first rest trial. The visual feedback (Fig. 1a) consisted of the normalised MEPs that were displayed as three bars representing the thumb, index, and little finger MEPs, respectively. Three white lines represented the baseline MEPs of the three finger muscles. If the bar exceeded the white line, the normalised MEP of the cued target finger was > 1, i.e., the current MEP was higher than the baseline MEP, indicating facilitation. If the bar was below the white line, the current MEP was below the baseline MEP (normalised MEP < 1), indicating suppression. If the bar of the cued target finger was both above the white bar and higher than the bars of the other two (non-target) finger muscles, the trial was deemed successful, and the bars were displayed in green. If not, the trial was deemed unsuccessful, and the bars were displayed in red. In a successful trial, participants could additionally reach up to three stars, one for each finger. To reach a star for the cued finger, the normalised MEP had to be > 150% of the other two non-target fingers. For the non-target fingers, the normalised MEPs had to be < 1.

### TMS-based neurofeedback training sessions

For the TMS-NF training sessions, we positioned the round coil over the vertex oriented to induce a posterior-anterior current flow in left M1. We first determined a stimulation intensity that elicited MEPs in all three finger muscles of the right hand. These MEPs should be in a range from which participants could up- and downregulate using motor imagery strategies, defined as 115% of the RMT of all three fingers. We therefore first measured the RMT of the three finger muscles, i.e., the minimum intensity needed to elicit MEPs of 50 μV amplitude with a probability of 0.5^84^ in *all* three finger muscles simultaneously at rest, using adaptive threshold hunting. Adaptive threshold hunting is based on maximum likelihood parameter estimation by sequential testing (PEST^85^) and was shown to be a highly reliable method to estimate the RMT with the advantage of using fewer trials compared to other methods^86,87^. PEST uses a probabilistic method to estimate the minimum TMS test stimulus (TS) intensity needed to elicit MEPs of a defined amplitude, here 50 μV for the RMT, in 50% of trials. We used an automated PEST script, implemented in MATLAB^88^, that incorporates the PEST function from the MTAT2.0 programme^89^ as described in^90^. The peak-to-peak amplitude of the MEP of the targeted muscle is calculated online and passed to the algorithm following pulse delivery. PEST then recommends a TS intensity for the following trial, which is more likely to be the RMT, based on whether the MEP amplitude reached the defined amplitude or not. We used a microcontroller to adjust the TS intensity automatically after each trial, prior to delivery of the next TMS pulse. This procedure was repeated for 20 trials to converge with sufficient confidence on an estimate of RMT^86^. As MEP amplitudes in the first trial are typically higher because of the novelty of the TMS sensation, we repeated the first trial, resulting in 21 trials for each block of adaptive threshold hunting.

We targeted the right APB, FDI, and ADM simultaneously, and therefore, the lowest amplitude of these three MEPs was passed to the PEST algorithm after each TMS pulse. As such, the resulting RMT was oriented to the finger muscle with the highest RMT. To ensure that the MEPs were not influenced by bgEMG, a trial was repeated automatically if the rms amplitude exceeded 10 μV in any of the three right-hand finger muscles. The experimenter visually controlled for a reliable convergence of the TS, i.e., a probability of approximately 0.5 to elicit MEPs of the defined amplitude in the last trials and otherwise repeated the RMT measure.

Following determination of RMT, we tested the estimated stimulation intensity for TMS-NF of 115% RMT and adjusted the intensity and / or the coil position if it did not elicit MEPs in all three finger muscles in each trial or if it resulted in ceiling effects in any of the three finger muscles. We then provided six blocks of TMS-NF in each training session. Each block consisted of 10 rest trials and 24 motor imagery trials, followed by a short break of 30 s between the blocks and a longer break after every second block. If the experimenter identified changes in corticospinal excitability based on MEP amplitudes during a session, the testing intensity was adjusted between blocks with longer breaks. During the first session, TMS-NF consisted of a blocked design, i.e., we cued a single finger for two consecutive blocks. This allowed participants to explore different motor imagery strategies. In the second session, we reduced the number of repetitions per finger to eight trials, and to four in the third session. The order of the blocks and cued fingers was pseudorandomised and balanced across participants. In the fourth session the trial order was completely interleaved and counterbalanced across cued fingers. An interleaved order of trials requires a change of the motor imagery strategies after each trial and, therefore, increases the difficulty. Studies have shown beneficial effects of such interleaved practice on delayed recall and long-term retention^91^. Mihelj et al.^31^ showed a high performance increase in a blocked trial order in TMS-NF. Thus, we designed a gradual change from a blocked to an interleaved order over sessions in this study. At the end of each session, participants noted down the strategies they had used for each of the fingers and rated each strategy on a scale from 1 (not successful at all) to 7 (very successful).

For the NF group, the post-training TMS session started with a short retraining consisting of two blocks of TMS-NF with four repetitions per finger.

For the feedback-free measures, we assessed two blocks that were identical to TMS-NF with an interleaved trial order, except that no visual feedback was provided. Instead, a white fixation cross appeared on the screen for the same duration (3s).

### Offline EMG data processing

Preprocessing of EMG data was performed using custom Python 3.7 scripts. EMG data from all six hand muscles were band-pass filtered (30-800 Hz) separately for the 5 – 105 ms of bgEMG before the TMS pulse was applied and for the 15 – 60 ms after the pulse that contained the MEP to avoid smearing of the MEP into the bgEMG. An additional 50 Hz notch filter was applied to the bgEMG data only. We calculated the rms of the bgEMG, the peak-to-peak MEP amplitude, and normalised the MEP and bgEMG of each motor imagery trial and finger muscle by the baseline of the rest trials in the corresponding TMS-NF block. We then split the dataset into training (NF 1 – 4) and feedback-free data. The training data is reported in the Supplementary Fig. 1a. Note that during TMS-NF, no online filters were applied. For all statistical analyses we used the feedback-free blocks from the pre- and post-training TMS sessions. For the three participants in the NF group that did not perform the feedback-free blocks in the post-training TMS session, we took the data from the feedback-free blocks in the fourth TMS-NF training session instead and showed that for the other 13 participants, motor imagery performance did not differ significantly in the fourth TMS-NF session vs post-training TMS-session (see Supplementary Fig. 1c).

During offline analysis we excluded all trials in which the rms amplitude of any of the muscles exceeded 7 µV (2.8 % of total feedback-free trials). We further excluded trials with rms values that were 2.5 SD above or below the mean bgEMG of each muscle (10.55 % of total feedback-free trials). Using the remaining trials, we quantified motor imagery performance, following similar procedures as in Mihelj et al.^31^ We calculated the MEP target ratio as the ratio between the normalised MEP of the cued target finger muscle and the higher of the non-target MEPs. An MEP target ratio > 1 indicates a finger-selective upregulation of corticospinal excitability; a value of 1 reflects no modulation; and values < 1 would show a finger-selective downregulation of corticospinal excitability. We then averaged the resulting MEP target ratio across all trials per participant and per session. We additionally computed the bgEMG target ratio using the bgEMG instead of MEPs and added it as a covariate in the linear mixed-effects model to control for subtle selective muscle contractions (bgEMG rms < 7 µV) in the motor imagery period.

### Paired-pulse TMS measurements

We used adaptive threshold hunting to assess short-interval intracortical inhibition (SICI), intracortical facilitation (ICF), and a single pulse (non-conditioned) protocol in the right FDI (i.e., index finger) while participants imagined moving either their index finger or while they imagined moving their thumb. This resulted in two motor imagery conditions where the index finger was either the target or a non-target finger.

We positioned the figure-of-eight-coil over the hotspot of the right FDI, i.e., the coil location eliciting the highest and most consistent MEPs in the right FDI. The coil was held tangential to the scalp at a 45° angle to the mid-sagittal line to achieve a posterior-anterior direction of current flow in the brain. This optimal coil location was registered in the neuronavigation software (Brainsight Frameless, Rogue Research Inc.). The position of the coil and the participant’s head were monitored in real-time using the Polaris Vicra Optical Tracking System (Northern Digital Inc.). First, we determined the RMT of the right FDI using adaptive threshold hunting (as described in **TMS-based neurofeedback training sessions**). Next, we measured the maximum MEP: We applied 10 pulses where the intensity of the first pulse was set to 50% of MSO, followed by three repetitions of 65%, 80%, and 95% of the MSO. The first trial was discarded because of the novelty of TMS sensation, and the maximum MEP was defined as the largest of the nine remaining MEPs without outliers.

For SICI and ICF we set the conditioning stimulus (CS) intensity to 70% RMT. The inter-stimulus interval (ISI) was set at 2 ms for SICI^42,66^ and 12 ms for ICF. In each block, we measured the TS during motor imagery which had a 50% probability of evoking an MEP of > 50% of the maximum MEP as target MEP. We tested one protocol per block, and two separate PEST protocols ran in an interleaved manner within a block to track the two TS of the motor imagery conditions (i.e., imagined index finger or imagined thumb movements) with 20 trials each. We determined the TS for both motor imagery conditions in the same block to control for changes in corticospinal excitability throughout the session. The cued finger (i.e., index or thumb) was repeated four times each. The structure of a trial was consistent with TMS-NF, except that a fixation cross and no feedback was presented for 2s after applying the TMS pulse(s). We applied a similar online bgEMG control as in TMS-NF, however, as we focused on motor imagery of the right index finger and thumb, the trial only paused when the bgEMG of the right APB or FDI exceeded 10 μV. For the other finger muscles, the dots representing the bgEMG turned yellow instead of red if bgEMG exceeded 10 μV and the trial proceeded normally. Participants were instructed to relax their muscles if a dot turned yellow but to primarily focus on motor imagery. If the bgEMG in the right APB or FDI exceeded 10 μV in the 5 – 105 ms before the CS (or TS in the single pulse protocol), the trial was repeated automatically. The order of stimulation protocols and which motor imagery condition was presented first in a block was balanced across participants but was kept consistent for the pre- and post-training sessions. The second assessed protocol was always the single pulse protocol. If the threshold of one of the two motor imagery conditions did not converge reliably, the block was repeated (see Supplementary Table 3 for number of repetitions per participant).

### Paired-pulse analysis

With the threshold hunting protocols, we determined the minimum stimulation intensity required to elicit an MEP of 50% of the maximum MEP amplitude in 50% of trials. We expressed inhibition (and facilitation) as the % change in intensity in the SICI (or ICF) protocol compared to the single pulse protocol. For inhibition, positive values indicate that a higher intensity was needed to elicit MEP amplitudes of at least the target MEP in the SICI compared to the single pulse protocol. For facilitation, positive values indicate that the ICF protocol resulted in a lower intensity than the single pulse protocol to elicited at least the target MEP amplitude.

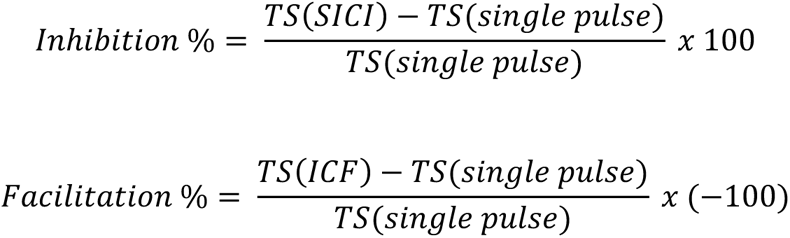

If a paired-pulse block was repeated, the plots of stimulation intensities and trials that showed positive and negative responses for each tested intensity were visually inspected by the experimenter and an independent, blinded researcher to decide which of the repetitions was used for further analysis: If possible, the thresholds for both motor imagery conditions (target vs non-target) were taken from the same block, unless the threshold of one motor imagery condition clearly converged better in another block. We then computed the pre- to post-training differences in inhibition, or facilitation, for the two motor imagery conditions.

#### fMRI tasks

We employed two paradigms in the pre- and post-training fMRI sessions to uncover neural changes after TMS-NF training. First, we assessed brain activity during imagined finger movements to analyse how finger-specific activity patterns change after TMS-NF training. To compare these activity patterns of imagined finger movements to those of executed movements, we additionally assessed motor execution in a paced finger-tapping task. Participants viewed a fixation cross centred on a screen through a mirror mounted to the head coil. For the motor imagery runs, participants were visually cued by the words ‘thumb’, ‘index’, ‘little’, or ‘rest’. Each motor imagery period was followed by a jittered rest period of 3 - 4 s during which a fixation cross was displayed instead of the task instruction. To ensure that participants did not execute any finger movements during this task, an experimenter visually controlled for finger movements inside the scanner room. If any movements were detected, we stopped the run, instructed the participant to refrain from executing finger movements, and repeated the run. We acquired four motor imagery runs using a blocked paradigm with block lengths of 7.5 s. In every run, each of the three fingers and rest were cued 12 times in a counterbalanced order, resulting in 48 trials per condition and session. Each motor imagery run lasted for 9 min 8 s.

During the motor execution runs, the participants’ right index, ring, middle and little fingers were placed on the buttons of a four-button response box, with the thumb placed on the side of the box. Participants viewed a fixation cross. They were then visually cued by the words ‘thumb’, ‘index’, ‘middle’, ‘ring’, ‘little’, or ‘rest’ appearing above the fixation cross to perform paced button presses with the corresponding finger (or to tap the side of the button box with the thumb) or to rest. The fixation cross blinked at 0.7 Hz to instruct the pace. In the rest condition, no fixation cross was displayed. We acquired six motor execution runs using a blocked paradigm with block lengths of 7.5 s. No breaks were provided between trials. In every run, each of the five fingers and rest were presented five times in a counterbalanced order, resulting in 30 trials per condition and session. Each motor execution run lasted for 4 min 5 s.

### fMRI data acquisition

We used a 3T Siemens Magnetom Prisma scanner with a 64-channel head-neck coil (Siemens Healthcare, Erlangen, Germany) to acquire fMRI data. For the anatomical T1-weighted images, we used a Magnetization Prepared Rapid Gradient Echo (MPRAGE) protocol with the following acquisition parameters: 160 sagittal slices, resolution = 1 x 1.1 x 1 mm^3^, field of view (FOV) = 240 x 240 x 160 mm, repetition time (TR) = 2300 ms, echo time (TE) = 2.25 ms, flip angle = 8°. For the task-fMRI data acquisition we used an echo-planar-imaging (EPI) sequence covering the whole brain and the cerebellum with the following acquisition parameters: 66 transversal slices, resolution = 2.2 mm^3^ isotropic, FOV = 210 x 210 x 145 mm, TR = 846 ms, TE = 30 ms, flip angle = 56°, acceleration factor = 6, and echo spacing = 0.6 ms. We acquired 636 and 278 volumes for each of the motor imagery and motor execution runs, respectively. To measure B0 deviations we used a fieldmap with the same resolution and slice angle as the EPI sequence and the following acquisition parameters: TR = 649 ms, TE1 = 4.92ms, TE2 = 7.38 ms.

### fMRI data preprocessing and co-registration

DICOM images were converted to nifti format using MRIcroGL v13.6 (https://www.nitrc.org/projects/mricrogl). MRI analysis was conducted using tools from FSL v.5.0.7 (http://fsl.fmrib. ox.ac.uk/fsl) unless stated otherwise. The following preprocessing steps were applied to the fMRI data using FSL’s Expert Analysis Tool (FEAT): motion correction using MCFLIRT^92^, brain extraction using the automated brain extraction tool (BET)^93^ , spatial smoothing using a 3 mm full-width at half-maximum (FWHM) Gaussian kernel, and high-pass temporal filtering with a 100 s cut-off. Non-brain tissue from the T1-weighted images of the pre- and post-training fMRI session was removed using BET and/or Advanced Normalization Tools (ANTs) v2.3.5 (http://stnava.github.io/ANTs) to receive a binarized mask of the extracted brain. Image co-registration was performed in separate, visually inspected steps. For each participant, we created a mid-space, i.e., an average space, between the T1-weighted images and its binarized brain masks of the pre and the post sessions. We then used the mid-space brain mask to brain extract the mid-space T1-weighted image. By using this T1-weighted mid-space for co-registration we ensured that the extent of reorientation required in the registration from functional to structural data was equal in the pre- and post-training fMRI sessions. Functional data were then aligned to the brain extracted T1-weighted mid-space, initially using six degrees of freedom and the mutual information cost function, and then optimised using boundary-based registration (BBR)^94^. To correct for B0 distortions, a fieldmap was constructed for B0 unwarping and added to the registration. For one participant, the fieldmap worsened co-registration in the MRI pre session and was therefore not applied. Three participants were taken out of the scanner for a brief break during the MRI pre-training session and the fieldmaps were only applied to the functional runs that were acquired with the same head position as the fieldmap. Structural images were transformed to Montreal Neurological Institute (MNI-152) standard space by nonlinear registration (FNIRT) with twelve degrees of freedom. The resulting warp fields were then applied to the functional statistical images.

Each functional run was assessed for excessive motion and excluded from further analyses if the absolute mean displacement was greater than half the voxel size (i.e., > 1.1 mm). This resulted in the exclusion of one motor execution fMRI run for two participants of the NF group.

### Univariate analysis

To assess univariate task-related activity of motor imagery and execution, time-series statistical analysis was carried out per run using FMRIB’s Improved Linear Model (FILM) with local autocorrelation, as implemented in FEAT. We defined one regressor of interest for each individual finger and obtained activity estimates using a general linear model (GLM) based on the gamma hemodynamic response function (HRF) and the temporal derivatives. We added nuisance regressors for the six motion parameters (rotation and translation along the x, y, and z-axis), as well as white matter (WM) and cerebrospinal fluid (CSF) time series.

For motor execution, we carefully inspected which finger participants used to press the button during each trial by examining the recorded button presses. When needed, we adjusted the finger movement regressors: If the button of a non-instructed finger was pressed during a motor execution trial, then we adjusted the regressors such that the trial was assigned to this non-instructed, moving, finger. If a button press indicated that the switch to the next cued finger was made with a delay, then we adjusted the corresponding block length and the movement onset of the next trial.

For motor imagery, we defined contrasts for each finger > rest, and overall task-related activity by contrasting all finger conditions > rest. We then averaged across runs at the individual participant level using fixed effect analysis. To define the motor imagery network, we entered the overall activity > rest contrast of the pre-training fMRI session of all participants (across the NF and control groups) into a mixed-effects higher-level analysis, and thresholded it at *Z* > 3.1, *p*_FWE_ < .05 at cluster level. Next, we aimed to test for activity changes from pre- to post-training and whether that differed between the groups. To do so, we defined pre > post and post > pre contrasts for the overall task-related activity at the individual participant level. We then used a mixed effect GLM to test for the group difference in a two-sample unpaired t-test. Additionally, to investigate group-specific effects in the pre- to post-training changes, we used mixed effect GLMs to compute one-sample t-tests on the pre > post and post > pre contrasts. Next, we investigated whether changes in the overall task-related activity were associated with changes in motor imagery performance (i.e., the MEP target ratio). To do so, we entered the pre- to post-training contrasts and the demeaned MEP target ratio changes in a mixed effect GLM to test the interaction effect, i.e., whether group differences in the pre- to post-training contrast maps vary as a function of motor imagery performance changes.

### Definition of regions of interest

We defined anatomical regions of interest (ROIs) based on the probabilistic Brodmann area (BA) parcellation using FreeSurfer v6.0 (https://surfer.nmr.mgh.harvard.edu/)^95–97^. We reconstructed the cortical surface of each individual participant’s T1-weighted mid-space image. We created a primary sensorimotor hand area ROI using similar procedures as in Kikkert et al.^3,98^. We first transformed BAs 1, 2, 3a, 3b, 4a, and 4p to volumetric space, merged them into an SM1 ROI, and filled any holes. Next, we non-linearly transformed axial slices spanning 2 cm medial/lateral to the anatomical hand knob on the 2 mm MNI standard brain (min-max MNI z-coordinates = 40 – 62) to each participant’s native structural space. Lastly, we used this mask to restrict the SM1 ROI and extracted an SM1 hand area ROI.

We further defined ROIs for dorsal and ventral premotor cortex (PMd and PMv), and supplementary motor area (SMA) by masking BA6 with the corresponding areas of the Human Motor Area Template (HMAT) atlas^100^ that were transformed into native space. For these masks, we then subtracted any overlap, as well as overlap with the SM1 hand area to avoid a voxel being assigned to multiple ROIs. Please see Supplementary Table 4 for the number of voxels of each ROI and participant.

### Representational similarity analysis (RSA)

While univariate analysis shows clusters of enhanced activity during imagined or executed finger movements, multivariate pattern analysis (MVPA) allows to investigate the fine-grained finger-specific activity patterns. Here, we used representational similarity analysis (RSA) to test the inter-finger distances of voxel-wise activity patterns elicited by individual finger motor imagery. We aimed to see whether these imagined finger movement representations became more distinct after TMS-NF training. To do so, we used the RSA toolbox^101^ and MATLAB R2015a. We computed the distance between the activity patterns for each finger pair in the SM1 hand ROI, SMA, PMd, and PMv using the cross-validated Mahalanobis distance, also called crossnobis distance^101^. Specifically, we extracted the voxel-wise parameter estimates (betas) for motor imagery of each finger > rest per run and the model fit residuals under an ROI. These extracted betas were then pre-whitened using the model fit residuals. To calculate the crossnobis distance for each finger pair, we used the four motor imagery runs as independent cross-validation folds and averaged the resulting distances across the folds. If it is impossible to statistically differentiate between motor imagery conditions (i.e. when this parameter is not represented in the ROI), the expected value of the distance estimate would be 0. If it is possible to distinguish between activity patterns, this value will be larger than 0^102^. We estimated the strength of the finger representation or ‘finger separability’ in each ROI as the average distance of all finger pairs.

A separability larger than 0 indicates that there is neural information content in the ROI that can statistically differentiate between motor imagery of individual fingers.

### Cross-condition classification

Next, we aimed to investigate whether neural activity patterns elicited by single-finger motor imagery became more similar to those observed during motor execution following TMS-NF training. To do so, we performed a cross-condition decoding analysis in the SM1 hand ROI, PMd, PMv, and SMA using the scikit-learn python library^103^ and nilearn^104^. We trained a classification algorithm to decode what finger was moved in each trial using the motor execution data. We then used this trained classifier to decode the motor imagery trials, i.e., which finger participants imagined moving. To create the training and test data, we computed single-trial parameter estimates using an HRF-based first-level GLM in SPM12 (http://www.fil.ion.ucl.ac.uk/spm/) using SPM’s default parameters. The design matrix consisted of individual regressors for each motor imagery and motor execution trial. This resulted in 48 parameter estimates per finger, session, and participant for motor imagery, and 30 for motor execution. Note that for motor execution, only thumb, index, and little finger trials were included. Ring and middle finger trials were modelled as regressors of no interest, as they were not analysed further for the present study. We added the same nuisance regressors as described in the univariate analysis section. Next, we extracted the voxel-wise parameter estimates below the specified SM1 hand ROI, SMA, PMd, and PMv, separately for each of these ROIs, trial, and participant. To ensure that a classifier can reliably decode executed finger movements, we first conducted a leave-one-run-out cross-validation within the motor execution condition using all runs of the pre- and post-training fMRI sessions, separately for each participant. For that, we scaled the data of the training data in a fold (i.e., eleven out of twelve runs) runs with the StandardScaler from the scikit-learn python library and trained a Support Vector Machine (SVM) with a linear kernel and default parameters of C = 1 and l2 regularization. We then applied the StandardScaler fitted on the eleven training runs on the left-out run and predicted the trials of this left-out run. We repeated this until each run once served as the left-out run. The classifier performance was based on the average classification accuracy from the cross-validation (Supplementary Fig. 3). To define the chance level, we generated a null distribution based on 1000 random permutations of the trial labels (i.e. ‘thumb’, ‘index’, ‘little’) for each participant. Then we computed an empirical *p*-value to evaluate the probability that the classification accuracy score was obtained by chance. For that, we divided the number of permutation-based classification accuracies that were greater than or equal to the true score +1, by the number of permutations + 1. To determine statistical significance at group level, we combined the empirical *p*-values of each participant for each ROI separately using Fisher’s method^105^.

For the cross-condition classification, we scaled the beta estimates across all runs of both the pre- and post-training sessions for each participant, but separately for the motor execution and imagery trials. Next, we trained an SVM with linear kernel and default parameters on all motor execution trials and tested it on all motor imagery trials, separately for the two sessions, to compare pre- to post-training decoding accuracy. To determine the empirical chance level, we shuffled the labels of the test set (i.e. motor imagery trials). We corrected the *p*-values for multiple comparisons within each group and ROI using the false discovery rate (FDR).

### Statistical analyses

Statistical analyses were performed in R v.4.3.1 (R Core Team, Vienna, Austria) and JASP v. 0.18.3 (JASP Team 2024, Netherlands). We used R packages lme4^106^ and lmertest^107^ to compute linear mixed-effects models. We defined Group (NF, control), Session (pre-training, post-training), or Motor imagery condition (target, non-target) as fixed effects and participant as a random effect. For each linear mixed-effects model, we evaluated the expected against observed residuals for normality and homoscedasticity using the R package DHARMA^108^ and did not find any violations. If the model revealed a significant interaction of the fixed effects, we computed post-hoc contrasts with the R package emmeans^109^. As we computed only one post-hoc contrast for each data set (i.e., each group), no correction for multiple comparisons was applied. For all other tests, we checked the data for violations against normality using the Shapiro-Wilk test. We then used standard classical parametric or non-parametric tests accordingly. We further used Bayesian tests (with default settings in JASP) to provide evidence for or against the null hypothesis and reported the Bayes factor BF_10_ following conventional cut-offs^110^.

Outliers were defined as > 2.5 SD from the group average. For the MEP target ratio, one participant of the NF group was classified as an outlier based on the TMS pre-training session. Removing this participant did not impact the conclusions of our statistical analysis (Supplementary Fig. 1b).

We used the R package effectsize^111^, to compute Cohen’s d based on F- and t-values from linear mixed-effects models and post-hoc contrasts, or we computed the effect sizes in JASP. Note that for negative t-values, we report effect sizes based on the absolute value. For Mann-Whitney tests, we report the rank biserial instead of Cohen’s d as effect size.

## Acknowledgments

We thank all participants of the study; S. Conticello, S. Leuenberger and J. Hajkova for assistance with piloting and data collection; L. Schönberg for assistance with data collection and preprocessing of fMRI data; W. Potok-Szybinska, X. Zhang and M. Altermatt for their guidance in the development of TMS protocols; D. Woolley for support with the TMS-NF software; E. Villar Ortega for the drawings; Ö. Özen for advice on the decoding analysis; and the Swiss Center for Musculoskeletal Imaging (SCMI) at Balgrist Campus for support with fMRI data acquisition. This project is supported by the Swiss National Science Foundation Grant 32003B_207719, the National Research Foundation, Prime Minister’s Office, Singapore under its Campus for Research Excellence and Technological Enterprise (CREATE) program (FHT), the AO Foundation and an ETH Zurich Research Grant. R.M. is supported by The Motor Neurone Disease Association UK (McMackin/Oct20/972-799), K.R. by the Health Research Board, Ireland, grant number EIA-2019-003, and S.K. by the Swiss National Science Foundation Ambizione Grant (PZ00P3_208996).

## Contributions

I.A.O., S.K. and N.W. conceptualised and designed the study. I.A.O, S.K., E.M., R.M. and K.R. programmed the task and analysis scripts. I.A.O., M.S.-L. and P.H. acquired the data. I.A.O., S.K. and N.W. planned the analysis. I.A.O. analysed the data. I.A.O., S.K and N.W. interpreted the data. I.A.O. drafted the manuscript and all authors substantively revised it.

