## Supplementary Material for "TMS-based neurofeedback training of mental finger individuation induces neuroplastic changes in the sensorimotor cortex"

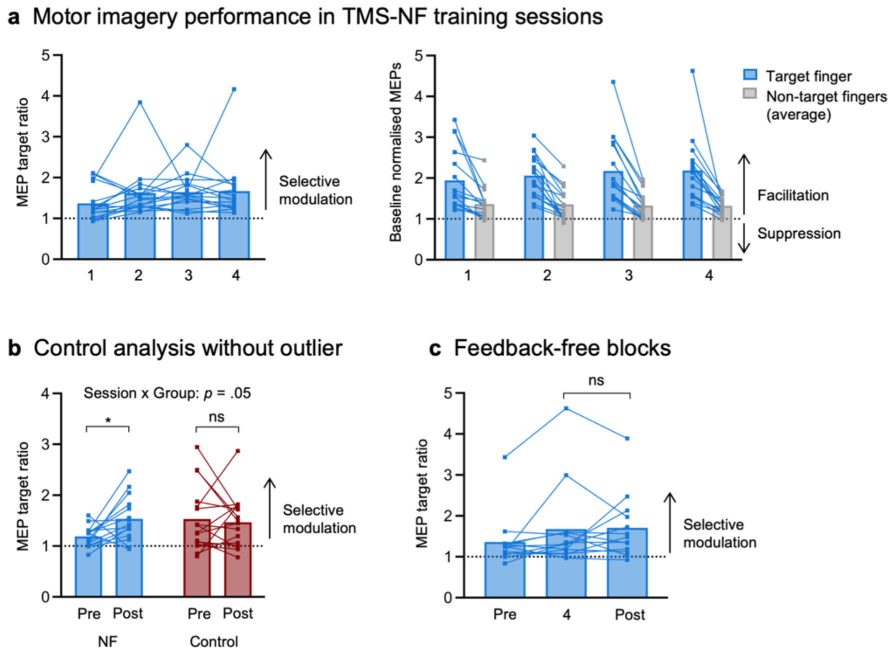

**Supplementary Figure 1. a)** TMS-NF training data. The x-axis corresponds to the four TMS-NF training sessions. Note that no statistics were applied on the training data as the motor imagery performance is confounded by increased difficulty through the transition from a blocked to an interleaved design. Left: MEP target ratio, i.e., the ratio between the normalised MEP (to the baseline at rest) of the target finger and the larger normalised MEP of the two non-target fingers. Values > 1 indicate a finger-selective modulation of corticospinal excitability. Right: Normalised MEPs of the target fingers (NF group = blue, control group = red) and the average normalised MEPs of the two non-target fingers (grey). **b)** MEP target ratio of the feedback-free blocks in the pre- and post-training sessions for both groups without the outlier participant in the NF group. The linear mixed-effects model revealed a trend for a Session (pre-training, post-training) by Group (NF, control) interaction ($F_{(1,30.10)}$ = 4.13, $p$ = .05, Cohen's $d$ = 0.74, 95% CI for Cohen's $d$: [0.00, 1.47]). Post-hoc contrasts show an increased MEP target from pre- to post-training for the NF group ($t_{(29.2)}$ = -2.38, $p$ = .02, Cohen's $d$ = 0.89, 95% CI for Cohen's $d$: [0.12, 1.66]) but not the control group ($t_{(28.9)}$ = .05, $p$ = .62, Cohen's $d$ = 0.18, 95% CI for Cohen's $d$: [-0.54, 0.90]). **c)** Feedback-free blocks of the NF group in the pre-training session, TMS-NF 4 session (i.e., last training session), and post-training session for n = 13 participants that underwent this measure to all three time points. A classical Wilcoxon signed-rank test showed no difference of TMS-NF 4 and the post-training session ($W$ = 46, $p$ = 1.00). Bayesian Wilcoxon signed-rank test revealed moderate evidence for the null hypothesis, i.e., no differences across the two measurement time points ($BF_{10}$ = 0.29). Squares depict data of individual participants. * p < .05; ns = non-significant.

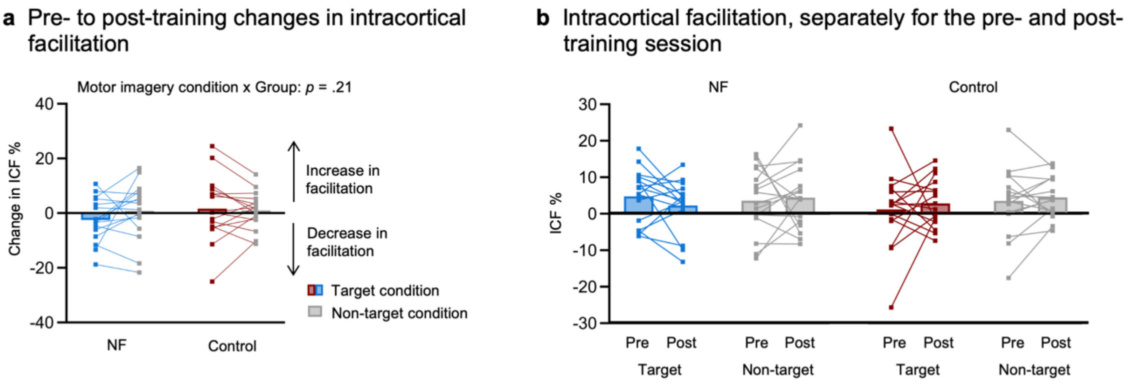

**Supplementary Figure 2**. No changes in intracortical facilitation (ICF) from pre- to post-training. **a)** Pre- to post-training changes in ICF. ICF was assessed with adaptive threshold hunting to determine the minimum testing stimulus intensity needed to elicit an MEP with an amplitude of at least 50% of the maximum MEP in 50% of trials. We measured ICF in the right index finger muscle during two motor imagery conditions: index as the target finger (motor imagery of index finger movements) vs index as an adjacent non-target finger (motor imagery of thumb movements). ICF is expressed as the % increase in the required testing stimulus intensity in the ICF protocol compared to a non-conditioned single pulse protocol during the same motor imagery condition. Positive scores indicate an increase in facilitation and negative scores indicate a decrease in facilitation after TMS-NF training. We did not observe any changes in facilitation from pre- to post-training for the target (blue for NF group; red for control group) relative to the non-target (grey) condition in any group (Group (NF, control) x motor imagery condition (target, non-target) interaction: $F_{(1,30)} = 1.64$, $p = 0.21$, Cohen's $d = 0.47$, 95% CI for Cohen's $d$: [-0.26, 1.19]). **b)** ICF for the pre- and post-training sessions separately. This data is shown for visualisation purposes only. Squares depict data of individual participants.

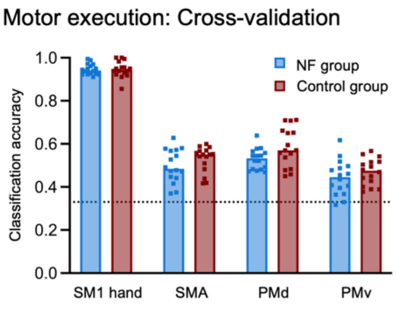

**Supplementary Figure 3**. Average classification accuracy from leave-one-run-out cross-validation within motor execution trials across the pre- and post-training sessions, separately for all ROIs and both groups (NF group = blue; control group = red). The dotted line represents the empirical chance level (33.33%). Classification accuracy for each ROI and group was significantly different from the empirical chance level (all $p < .0001$). Squares depict data of individual participants.

**Supplementary Table 1a.** Activation clusters, corresponding size, anatomical region, FW-corrected p-value for multiple comparisons, peak coordinate in MNI space, and maximum z-value of the reported contrast of motor imagery (pre-training session, across all fingers and groups) vs rest thresholded at z > 3.1. Reported anatomical labels were determined using the Jülich Histological[1], the Harvard-Oxford cortical[2] and subcortical structural[3], and the probabilistic cerebellar atlases[4], correspond to the location of maxima within each cluster.

| **Both groups, pre-training session: Motor imagery > rest** | | | | | | | |
|---|---|---|---|---|---|---|---|
| Cluster | # voxels | Region of peak | $p_{(FWE)}$ | Peak coordinates | | | z-value |
| | | | | X | Y | Z | |
| 1 | 6300 | Left premotor cortex | 0 | -56 | 6 | 20 | 6.5 |
| 2 | 1947 | Left inferior parietal lobe | 1.4e-42 | -50 | -28 | 36 | 5.99 |
| 3 | 1161 | Right premotor cortex | 8.24e-30 | 56 | 6 | 18 | 5.77 |
| 4 | 922 | Right VI | 2.17e-25 | 34 | -58 | -24 | 6.26 |
| 5 | 649 | Right VIIIa | 7.77e-20 | 28 | -60 | -54 | 6.54 |
| 6 | 134 | Left visual cortex V2 | 3.93e-06 | -12 | -92 | -4 | 5.21 |
| 7 | 129 | Right visual cortex V1 | 5.96e-06 | 14 | -90 | 0 | 4.7 |
| 8 | 115 | Right premotor cortex | 2.02e-05 | 26 | -12 | 50 | 4.15 |
| 9 | 103 | Left middle frontal gyrus | 5.97e-05 | -40 | 38 | 28 | 4.17 |
| 10 | 95 | Left VI | 0.000126 | -30 | -54 | -28 | 4.53 |
| 11 | 85 | Right hippocampus | 0.00033 | 20 | -14 | -18 | 5.31 |
| 12 | 72 | Left VIIIa | 0.00122 | -30 | -58 | -56 | 5 |
| 13 | 68 | Left Crus I | 0.00186 | -44 | -56 | -30 | 5.41 |
| 14 | 63 | Left hippocampus | 0.00317 | -20 | -16 | -14 | 5.2 |
| 15 | 46 | Left thalamus | 0.0218 | -8 | -20 | -2 | 5.43 |
| 16 | 40 | Right anterior intra-parietal sulcus/ superior parietal lobule | 0.045 | 36 | -34 | 40 | 3.89 |

**Supplementary Table 1b.** Activation clusters, corresponding size, anatomical region, FW-corrected p-value for multiple comparisons, peak coordinate in MNI space, and maximum z-value of the reported pre- to post-training contrasts thresholded at z > 3.1. Reported anatomical labels were determined using the Jülich Histological[1], the Harvard-Oxford cortical[2] and subcortical structural[3], and the probabilistic cerebellar atlases[4], correspond to the location of maxima within each cluster.

| **NF group: Pre-training > Post-training** | | | | | | | |
|---|---|---|---|---|---|---|---|
| *None* | | | | | | | |
| **NF group: Post-training > Pre-training** | | | | | | | |
| Cluster | # voxels | Region of peak | $p_{(FWE)}$ | Peak coordinates | | | z-value |
| | | | | X | Y | Z | |
| 1 | 61 | Right precuneous | 0.00115 | 10 | -70 | 36 | 3.76 |
| **Control group: Pre-training > Post-training** | | | | | | | |
| *None* | | | | | | | |
| **Control group: Post-training > Pre-training** | | | | | | | |
| 1 | 380 | Right inferior parietal lobule | 9.96e-17 | 56 | -44 | 26 | 4.33 |
| 2 | 190 | Right anterior intra-parietal sulcus | 4.92e-10 | 38 | -52 | 38 | 4.23 |
| 3 | 162 | Right middle temporal gyrus | 6.94e-09 | 60 | -20 | -10 | 4.15 |
| 4 | 107 | Bilateral posterior cingulate gyrus / precuneous | 2.09e-06 | 2 | -48 | 36 | 3.88 |
| 5 | 70 | Right middle frontal gyrus | 0.00017 | 28 | 18 | 54 | 3.92 |
| 6 | 63 | Left superior lateral occipital cortex | 0.000425 | -30 | -74 | 26 | 4.43 |
| 7 | 61 | Left Crus I | 0.000556 | -12 | -76 | -30 | 4.12 |
| 8 | 42 | Right superior parietal lobule | 0.00832 | 4 | -38 | 48 | 4.32 |
| 9 | 41 | Right middle frontal gyrus | 0.00969 | 50 | 20 | 30 | 3.68 |

**Supplementary Table 1c.** Activation clusters, corresponding size, anatomical region, FW-corrected p-value for multiple comparisons, peak coordinate in MNI space, and maximum z-value of the reported interaction analysis where the relationship between pre- to post-training changes and motor imagery performance changes differs between the NF and control groups, thresholded at z > 3.1. Reported anatomical labels were determined using the Jülich Histological[1], the Harvard-Oxford cortical[2] and subcortical structural[3], and the probabilistic cerebellar atlases[4], correspond to the location of maxima within each cluster.

| **Interaction effect of Group and Covariate: NF group > Control group** | | | | | | | |
|---|---|---|---|---|---|---|---|
| Cluster | # voxels | Region of peak | $p_{(FWE)}$ | Peak coordinates | | | z-value |
| | | | | X | Y | Z | |
| 1 | 229 | Left primary motor cortex | 7.59e-11 | -34 | -28 | 58 | 5.01 |
| 2 | 132 | Left frontal pole | 4.17e-07 | -34 | 52 | 4 | 4.08 |
| 3 | 128 | Right V | 5.96e-07 | 20 | -44 | -20 | 4.62 |
| 4 | 128 | Right paracingulate gyrus | 5.69e-07 | 12 | 14 | 38 | 4.59 |
| 5 | 100 | Right SMA | 1.1e-05 | -4 | 2 | 60 | 4.69 |
| 6 | 70 | Right VIIIa | 0.000331 | 14 | -68 | -48 | 4.58 |
| 7 | 66 | Left temporal pole | 0.00054 | -50 | 16 | -8 | 4.55 |
| 8 | 60 | Left premotor cortex | 0.00115 | -52 | -2 | 34 | 4.14 |
| 9 | 53 | Left precuneous | 0.00284 | -8 | -72 | 28 | 4.12 |
| 10 | 45 | Left primary somatosensory cortex | 0.00844 | -50 | -36 | 56 | 4.12 |
| 11 | 38 | Left precuneous | 0.023 | -8 | -72 | 46 | 4.6 |
| 12 | 37 | Left superior parietal lobule | 0.0266 | -22 | -66 | 54 | 4 |
| 13 | 33 | Left premotor cortex | 0.0484 | -26 | -6 | 50 | 3.87 |
| **Interaction effect of Group and Covariate: Control group > NF group** | | | | | | | |
| 1 | 37 | Left middle temporal gyrus | 0.0266 | -64 | -6 | -28 | 4.48 |

**Supplementary Table 2a.** Verbatim instructions for motor imagery tasks. These instructions were provided at the beginning of the first session (TMS pre-training session) where participants did not receive any feedback yet and were identical for the NF group and control group.

| |
|---|
| • Increase activity by **vividly imagining to move** (**feeling** not seeing!) **the instructed finger** of the right hand, for example by imagining:<br> • to move the finger up/down, left/right.<br> • to press a button, playing piano, typing on a keyboard.<br> • pulses in the muscle.<br> • …<br>• Complex or forceful movements will give more activation than weak or simple movements.<br>• The imagined movement should **only involve the instructed finger** and no others. |
| • **Decrease activity for the not instructed fingers**, for example by imagining:<br> • that the other fingers were very cold, as if in a bucket of ice water.<br> • would not belong to the body or not exist at all.<br> • … |
| • Imagine as long as you can see the instruction on the screen. There will be one or two TMS pulse(s) during this time.<br>• Ensure no actual muscle activity anywhere in the body. Keep also your face muscles completely relaxed. |

**Supplementary Table 2b**. Self-reported strategies used during TMS-NF by participants in the NF group ($n$ = 16) and number of participants that have reported these strategies, separately for each finger. Please note that participants were allowed to use multiple strategies. All reported strategies involved motor imagery. The strategies in the upper block involve additional imagined sensory feedback by touch or pressure, while the strategies in the bottom block include rather imagined proprioceptive or thermoceptive feedback.

| **Strategy** | **Thumb** | **Index** | **Little** |
|---|---|---|---|
| imagine to tap or press on button, keyboard/piano key, phone, pillow | 11 | 15 | 10 |
| imagine to move the finger over pillow | 1 | 3 | 1 |
| imagine to pull or plug a string/object (e.g. violin, open a soda can), digging in sand, pinky promise | 2 | 1 | 2 |
| imagine that whole bodyweight is hold by this finger | 0 | 0 | 1 |
| imagine that finger sticked in a (hot or cold) rocky hole, focus on climbing texture | 1 | 1 | 0 |
| imagine to hold, bend or move the fingers to the left, right, up, downwards, in circles (e.g. thumbs up, point on something, say "no" with index finger, dancing with the finger) | 12 | 11 | 12 |
| imagine that other fingers are not here, only focus on the target finger | 1 | 1 | 1 |
| imagine that finger is heavy/energized and other fingers are light | 1 | 1 | 0 |
| imagine that finger get warm and other fingers get cold<br>OR just the target finger get cold/warm | 2 | 1 | 2 |
| imagine the muscle contraction in the finger | 1 | 0 | 1 |

**Supplementary Table 2c**. Self-reported strategies during the fMRI pre- and post-training session, separately for each participant, sorted for the NF and control group. Please note that we only started to collect self-report of strategies in the fMRI session after study onset (for NF group from participant nr. 9, and for control group from participant nr. 4 on). For participants in the NF group, the strategies from the post-training session corresponds to the strategies used in the TMS-NF training.

| NF group | Session | Thumb | Index | Little |
|---|---|---|---|---|
| 9 | pre | dancing with the finger | dancing with the finger | dancing with the finger |
| | post | feeling the muscle contraction | pressing the index finger against the pillow | dancing with the finger |
| 10 | pre | thumbs up | pointing on something | keyboard |
| | post | thumbs up<br>partying of the thumb<br>pushing something with the finger | pointing on something<br>partying of the index finger | pushing button<br>partying of the little finger<br>moving the finger towards left and right |
| 11 | pre | feeling the thumb close<br>opening a can of beer with the thumb | flicking a paper ball<br>pointing at someone<br>opening a can of beer<br>rubbing the table | stretching the finger<br>closing the finger |
| | post | pushing thumb outwards (in and out) in little movements | pushing index finger outwards | pushing little finger outwards |
| 12 | pre | movements in the joint<br>movement of the skin due to the movements in all directions | movements in the joint<br>movement of the skin due to the movements in all directions<br>-> especially flexion | movements in the joint<br>movement of the skin due to the movements in all directions<br>-> especially flexion |
| | post | pressing space bar, exaggerated extension | pressing key on keyboard | pressing shift on keyboard |
| 13 | pre | thumb stuck and wriggling out<br>frozen/paralyzed other fingers<br>moving thumb | finger stuck and wriggling out | sipping a cup of tea like a gentleman |
| | post | sticking into rocky holes - something hot and cold | move heavily on the climbing, texture feeling | negative incidents involving little finger, little finger injuries |
| 14 | pre | pushing button inwards | pushing button downwards | pressure downwards |
| | post | pressure towards left downwards | pressure towards left downwards | pressure downwards |
| 15 | pre | moving up- and downwards | making circles, up- and downward, left and right | scratching on surface |
| | post | Lighter | pressing | spreading |
| 16 | pre | pushing elevator button | pushing elevator button | pushing elevator button |
| | post | pressing down | pressing down | pressing down |
| **Control group** | **Session** | **Thumb** | **Index** | **Little** |
| 4 | pre | pushing palm | pushing button | *did not recall* |
| | post | pushing palm | brooding cheese | pushing button |
| 5 | pre | "thumbs up" multiple time | holding up index finger | holding up little finger |
| | post | typing | typing | typing |
| 6 | pre | moving thumb up- and downwards, | moving index up- and downwards, | moving little finger up- and downwards, |

| | | | | |
|---|---|---|---|---|
| | | pushing piano key,<br>typing,<br>moving thumb over surface | pushing piano key,<br>typing,<br>moving index over surface | pushing piano key,<br>typing,<br>moving little finger over surface |
| | post | moving thumb up- and downwards,<br>pushing piano key,<br>typing,<br>moving thumb over surface | moving index up- and downwards,<br>pushing piano key,<br>typing,<br>moving index over surface | moving little finger up- and downwards,<br>pushing piano key,<br>typing,<br>moving little finger over surface |
| 7 | pre | playing musical instrument,<br>scratching over table,<br>moving up and down in the air | playing musical instrument,<br>scratching over table,<br>moving up and down in the air | playing musical instrument,<br>scratching over table,<br>moving up and down in the air |
| | post | moving it in a round circle,<br>cutting the cake with one finger | moving it in a round circle,<br>cutting the cake with one finger | moving it in a round circle,<br>cutting the cake with one finger |
| 8 | pre | moving finger to the left and right,<br>spreading | typing | moving finger to the left and right,<br>spreading |
| | post | bending | typing | spreading |
| 9 | pre | kneading bread,<br>testing if it is ready to bake,<br>making circles | looking through a map,<br>pointing at landmark/location,<br>making circles | making circles,<br>imagining the circles and other fingers in ice |
| | post | pressing my side,<br>making circles,<br>testing bread dough | looking through a map,<br>pointing at landmark,<br>making circles and lifting | lifting the little finger |
| 10 | pre | counting number of guests with counting device,<br>welcoming passengers | set timer on the oven | Groove of MRI |
| | post | home button on phone,<br>counting number of guests | set timer on the oven | typing on the edge of MRI |
| 11 | pre | pressing a button | pressing a button | moving finger up and down |
| | post | pressing a button | pressing a button | moving finger up and down<br>pressing a button (as alternative) |
| 12 | pre | pressing on surface | pressing on surface | stretching outwards |
| | post | pressing outwards | pressing on surface | stretching outwards |
| 13 | pre | thumb game,<br>twiddling the thumbs,<br>à in sign language | 'worm song' with finger,<br>making circles with index finger | 'worm song' with finger,<br>making circles with little finger |
| | post | à in sign language | 'worm song' | 'worm song' |
| 14 | pre | thumb war,<br>pressing controller button | pressing controller button,<br>rotating a ring around using index finger,<br>tapping on the desk | pinky swear,<br>moving little finger around into a water bowl |
| | post | extending my thumb<br>thumb war,<br>tapping on smartphone | sliding finger on surface,<br>especially paper | pinky swear |
| 15 | pre | rotating controller of a beamer, | rotating controller of a beamer, | movements of extension, adduction, abduction, |

| | | | | |
|---|---|---|---|---|
| | | movements of pressing, extension, adduction, abduction and rotation | movements of pressing, extension, adduction, abduction and rotation | rotation, pulling a hook downwards |
| | post | rotating controller of a beamer,<br>movements of pressing, extension, adduction, abduction and rotation,<br>pressing spacebar on keyboard,<br>using gaming controller | rotating controller of a beamer,<br>movements of pressing, extension, adduction, abduction and rotation,<br>pressing piano key and keyboard | movements of extension, adduction, abduction, rotation, pulling a hook downwards,<br>pressing piano key |
| 16 | pre | making circles (like holding a joystick),<br>up- and downwards movements,<br>grabbing a handle | pulling a trigger,<br>picking nose,<br>scratching the head | moving finger left, right, up, down, pushing outwards,<br>holding a rope, |
| | post | moving finger in figure of 8, up and down,<br>apply sunscreen | Tickling,<br>swiping a screen,<br>moving finger in figure of 8 | moving finger to the side,<br>swipe inside of a ham jar |

**Supplementary Table 3.** Number of repetitions for the paired-pulse TMS protocols in the pre- and post-training TMS sessions.

| **NF group** | **Session** | **SICI** | **ICF** | **Single pulse** |
|---|---|---|---|---|
| 1 | pre | 1 | 1 | 1 |
| | post | 1 | 2 | 1 |
| 2 | pre | 2 | 1 | 1 |
| | post | 1 | 1 | 2 |
| 3 | pre | 1 | 1 | 2 |
| | post | 1 | 3 | 1 |
| 4 | pre | 1 | 1 | 2 |
| | post | 2 | 1 | 2 |
| 5 | pre | 2 | 1 | 2 |
| | post | 1 | 1 | 2 |
| 6 | pre | 2 | 1 | 2 |
| | post | 1 | 1 | 2 |
| 7 | pre | 2 | 1 | 1 |
| | post | 2 | 1 | 2 |
| 8 | pre | 2 | 1 | 2 |
| | post | 1 | 1 | 2 |
| 9 | pre | 2 | 1 | 1 |
| | post | 1 | 2 | 1 |
| 10 | pre | 2 | 1 | 1 |
| | post | 1 | 1 | 2 |
| 11 | pre | 1 | 1 | 2 |
| | post | 2 | 1 | 1 |
| 12 | pre | 2 | 2 | 1 |
| | post | 2 | 1 | 1 |
| 13 | pre | 1 | 1 | 2 |
| | post | 1 | 1 | 2 |
| 14 | pre | 1 | 2 | 2 |
| | post | 2 | 1 | 1 |
| 15 | pre | 2 | 1 | 2 |
| | post | 1 | 2 | 1 |
| 16 | pre | 1 | 2 | 2 |
| | post | 2 | 1 | 1 |
| **NF group** | **Session** | **SICI** | **ICF** | **Single pulse** |
| 1 | pre | 1 | 1 | 2 |
| | post | 1 | 2 | 1 |
| 2 | pre | 2 | 1 | 2 |
| | post | 2 | 2 | 2 |
| 3 | pre | 1 | 1 | 2 |
| | post | 1 | 1 | 2 |
| 4 | pre | 2 | 1 | 1 |
| | post | 1 | 1 | 2 |
| 5 | pre | 2 | 1 | 2 |
| | post | 1 | 1 | 1 |
| 6 | pre | 1 | 2 | 2 |
| | post | 1 | 1 | 2 |
| 7 | pre | 1 | 1 | 2 |
| | post | 1 | 2 | 1 |
| 8 | pre | 1 | 1 | 1 |
| | post | 1 | 1 | 2 |
| 9 | pre | 1 | 2 | 2 |
| | post | 2 | 1 | 1 |
| 10 | pre | 1 | 1 | 2 |
| | post | 1 | 1 | 2 |

| | | | | |
|---|---|---|---|---|
| 11 | pre | 1 | 1 | 2 |
| | post | 2 | 1 | 1 |
| 12 | pre | 1 | 2 | 2 |
| | post | 1 | 1 | 2 |
| 13 | pre | 1 | 2 | 1 |
| | post | 1 | 2 | 2 |
| 14 | pre | 2 | 1 | 1 |
| | post | 1 | 2 | 1 |
| 15 | pre | 2 | 1 | 1 |
| | post | 1 | 1 | 2 |
| 16 | pre | 2 | 1 | 1 |
| | post | 1 | 2 | 1 |

**Supplementary Table 4.** Number of voxels for all ROIs in individual participant's space.

| NF group | SM1 hand | SMA | PMd | PMv |
|---|---|---|---|---|
| 1 | 1377 | 533 | 1083 | 213 |
| 2 | 1449 | 565 | 906 | 351 |
| 3 | 1088 | 358 | 686 | 214 |
| 4 | 1483 | 529 | 746 | 171 |
| 5 | 1761 | 718 | 576 | 224 |
| 6 | 1244 | 479 | 956 | 312 |
| 7 | 1645 | 450 | 944 | 227 |
| 8 | 1115 | 420 | 658 | 190 |
| 9 | 1436 | 460 | 965 | 168 |
| 10 | 1183 | 536 | 885 | 321 |
| 11 | 1685 | 932 | 942 | 255 |
| 12 | 1464 | 742 | 884 | 206 |
| 13 | 1366 | 507 | 1017 | 245 |
| 14 | 1514 | 381 | 1053 | 231 |
| 15 | 2021 | 575 | 1026 | 199 |
| 16 | 1223 | 520 | 695 | 241 |
| **Control group** | **SM1 hand** | **SMA** | **PMd** | **PMv** |
| 1 | 1407 | 527 | 1051 | 203 |
| 2 | 1222 | 340 | 802 | 213 |
| 3 | 1959 | 877 | 888 | 290 |
| 4 | 1312 | 606 | 935 | 291 |
| 5 | 1322 | 588 | 1064 | 222 |
| 6 | 1546 | 705 | 1102 | 330 |
| 7 | 1293 | 641 | 638 | 164 |
| 8 | 1427 | 486 | 647 | 209 |
| 9 | 1354 | 544 | 994 | 216 |
| 10 | 1359 | 777 | 783 | 167 |
| 11 | 1336 | 483 | 939 | 222 |
| 12 | 1418 | 564 | 1019 | 291 |
| 13 | 1268 | 402 | 960 | 194 |
| 14 | 1290 | 614 | 688 | 227 |
| 15 | 1471 | 910 | 516 | 236 |
| 16 | 1874 | 641 | 1055 | 240 |
